# Atg8-LC3 controls systemic nutrient surplus signaling from flies to humans

**DOI:** 10.1101/2023.05.31.543119

**Authors:** Aditi Madan, Kevin P. Kelly, Camille E. Sullivan, Michelle E. Poling, Ava E. Brent, Mroj Alassaf, Julien Dubrulle, Akhila Rajan

**Author notes:** Present address: Department of Biological Sciences, Columbia University, New York, New York, USA. Present address: ^ Brandeis University, Waltham, Massachusetts, USA. equal contribution (Aditi Madan, Kevin P. Kelly).

## Abstract

Organisms experience constant nutritional flux, and homeostatic mechanisms evolved to operate at the nexus of extreme nutritional states - scarcity and surplus. Thus, we surmised that decoding bidirectional molecular switches that operate at the interface of scarcity response and surplus signaling will enable the development of strategies to treat disorders that arise from nutrient imbalance states. Adipocytes secrete leptin, an interleukin protein, which signals nutrient surplus to the central brain to regulate feeding and energy expenditure. We report that Atg8-LC3-family proteins, best known for their role in autophagy, are required for leptin secretion in *Drosophila* and human adipocytes. Atg8-LC3 genetic knockdown and point mutations to the Atg8-LC3 interaction motif (AIM/LIR) of leptin, and its functional ortholog in *Drosophila*, Upd2, disrupt adipokine secretion and increase adipokine retention in human and fly cells. At an organismal level, Atg8-driven Upd2 retention increases organismal resilience to nutrient extremes by rewiring the transcriptome, organismal feeding behavior, and hunger response. Comparative proteomic analyses reveal that LC3 directs leptin to an exosome secretory pathway. We use genetic knockdown in primary human adipocytes to establish that LC3 is required for leptin secretion in a physiologically relevant mammalian system. Hence, we uncover a previously unknown and evolutionarily conserved role for Atg8-LC3 in promoting adipocyte-brain nutrient surplus signaling. We propose that Atg8-LC3’s bidirectional role in nutrient sensing-conveying nutrient surplus and responding to nutrient deprivation-enables organisms to manage nutrient flux effectively.

## Introduction

Organisms constantly evaluate their basal nutritional reserves to decide whether to allocate resources to expensive functions that enhance fitness or conserve energy. Adipokines, secreted by fat cells, provide systemic information on nutrient-reserve information and signal the nutrient surplus state. Leptin in mammals^1^ and its functional ortholog in fruit flies Unpaired2 (Upd2)^2^ are primary adipokines released in proportion to fat stores^3-5^. They impinge on brain circuits that control energy expenditure, appetite, and overall metabolism^2,6^. In surplus nutrient states, adipokines convey a permissive signal, indicating that energy resources can be devoted to costly activities like immunity and reproduction. During periods of scarcity, their secretion is reduced, signaling an energy deficit and enabling organisms to conserve energy.

In humans^3,4^ and mice^1,7^, starvation reduces leptin levels, which is crucial for a neuroendocrine response to reduced energy reserves. Exogenously injecting leptin into starved mice dysregulates neuroendocrine physiology^1^. High levels of circulating leptin, termed ‘hyperleptinemia’, lead to ineffective leptin signaling to the brain^8,9^. Therefore, pinpointing regulators of leptin secretion might provide novel avenues to improve dysfunctional energy balance by restoring adipocyte-brain signaling. Upd2 is a JAK/STAT pathway agonist^10^ and a functional ortholog of the human adipokine leptin^2^. In previous studies, we identified that human leptin could functionally substitute and rescue metabolic defects of a *Upd2Δ* loss-of-function allele^2^. Given this evolutionary conservation, we used the fly system as a discovery tool to identify acute, cell-intrinsic mechanisms that regulate adipokine secretion.

Organisms experience constant nutritional flux. Hence homeostatic mechanisms evolved to operate at this nexus state between two extreme nutritional states - scarcity and surplus. We surmised that an overlapping set of molecules that allow organisms to sense and respond to these two mutually exclusive states-scarcity and surplus must exist to respond to nutrient flux adeptly. Therefore, we explored whether proteins pivotal to nutrient deprivation regulate adipokine secretion.

The nutrient-scavenging pathway autophagy is a primary mechanism by which organisms adapt to metabolic stress^11,12^. In adipocytes, autophagy controls lipid metabolism through lipophagy, and chaperone-mediated autophagy is critical for adipocyte differentiation^13-15^. Central players in the autophagy pathway are ubiquitin-like protein family members^16^, LC3/GABARAP in mammals, and Atg8 in flies and yeast^17^. The Atg8/LC3 ubiquitin-like protein is a primary component of the autophagy pathway^17^. When conjugated to the lipid moiety phosphatidylethanolamine (PE)^18^, Atg8 regulates the fusion of autophagosomes to lysosomes, resulting in cargo degradation^19^. In addition to degradative roles, in a process termed atg8lyation, mammalian Atg8s (LC3s) are conjugated to single membranes to control diverse cellular processes from phagocytosis to secretion^20-24^. These include but are not limited to LC3-associated phagocytosis (LAP)^25^, viral replication^26^, extracellular vesicle secretion (LDELS)^27^, and LC3-associated endocytosis (LANDO)^28^. It has now emerged that conjugation of Atg8 on single membranes (CASM) to both PE and PS lipids allows Atg8 to participate in a noncanonical autophagic process^29,30^. Atg8 and other autophagy (*ATG*) genes participate in autophagy-independent trafficking processes^21^, for example, in unconventional secretion of yeast Acb1^22^ and mammalian interleukins ^23,24^.

Given LC3’s role in nutrient scavenging^17^ and its documented role in secretion^22-24,27,31^, we specifically examined whether Atg8-LC3-family proteins control the secretion of *Drosophila* Upd2 and human leptin from adipocytes. Note we will refer to Atg8 in the context of *Drosophila* experiments and LC3 when describing mammalian data. By testing the role of Atg8 and LC3 in various cell types *in vitro*, in *Drosophila* adipocytes *in vivo*, and in primary human subcutaneous adipocytes, we conclude that LC3 family proteins promote leptin secretion. Comparative proteomics analyses of human cells expressing secretion-competent versus -incompetent leptin revealed that leptin is secreted via LC3-mediated exosome secretion. Altogether, we identify a previously underappreciated role for noncanonical functions of Atg8-LC3 proteins in signaling a nutrient surplus state from adipocytes to the brain. The role of Atg8-LC3 protein in nutrient deprivation has been long appreciated ^32^. Now we identify a central role for Atg8-LC3 in promoting the secretion of hormones that signal nutrient surplus. Based on our work, we propose that Atg8-LC3 plays a bidirectional role in nutrient response-promoting nutrient surplus signaling via adipokine release and orchestrating autophagy^17^ in response to starvation.

## Results

### Atg8 regulates Upd2 secretion in *Drosophila* cells

We set out to address whether Upd2 release from cells is coupled to nutrient state using *Drosophila* S2R+ cells as a starting point. Previously, functional genomics studies have utilized the *Drosophila* S2R+ system to provide important insights into conserved pathways of secretion^33^, lipid droplet formation^34^, and nutrient sensing^35,36^. Since S2R+ cells respond robustly to amino-acid (AA) deprivation^35,36^, we can test the effect of acute nutrient deprivation on Upd2 intracellular retention versus secretion.

We transiently transfected cells with Upd2-WT::GFP (Wild-type [WT] Upd2 C-terminally tagged with GFP) and assayed GFP localization in cells cultured in complete media (fed) or in amino-acid (AA)-depleted media (starved). We observed a significant increase in Upd2 accumulation within 8 hours of AA deprivation, with the maximum accumulation observed within 4-12 hours (**Fig. 1A, A’, S1**). Refeeding the cells with complete media reversed the intracellular accumulation significantly (**Fig. 1A, A’**), highlighting the acute impact of the nutrient state on Upd2’s intracellular retention.

**Figure 1.**
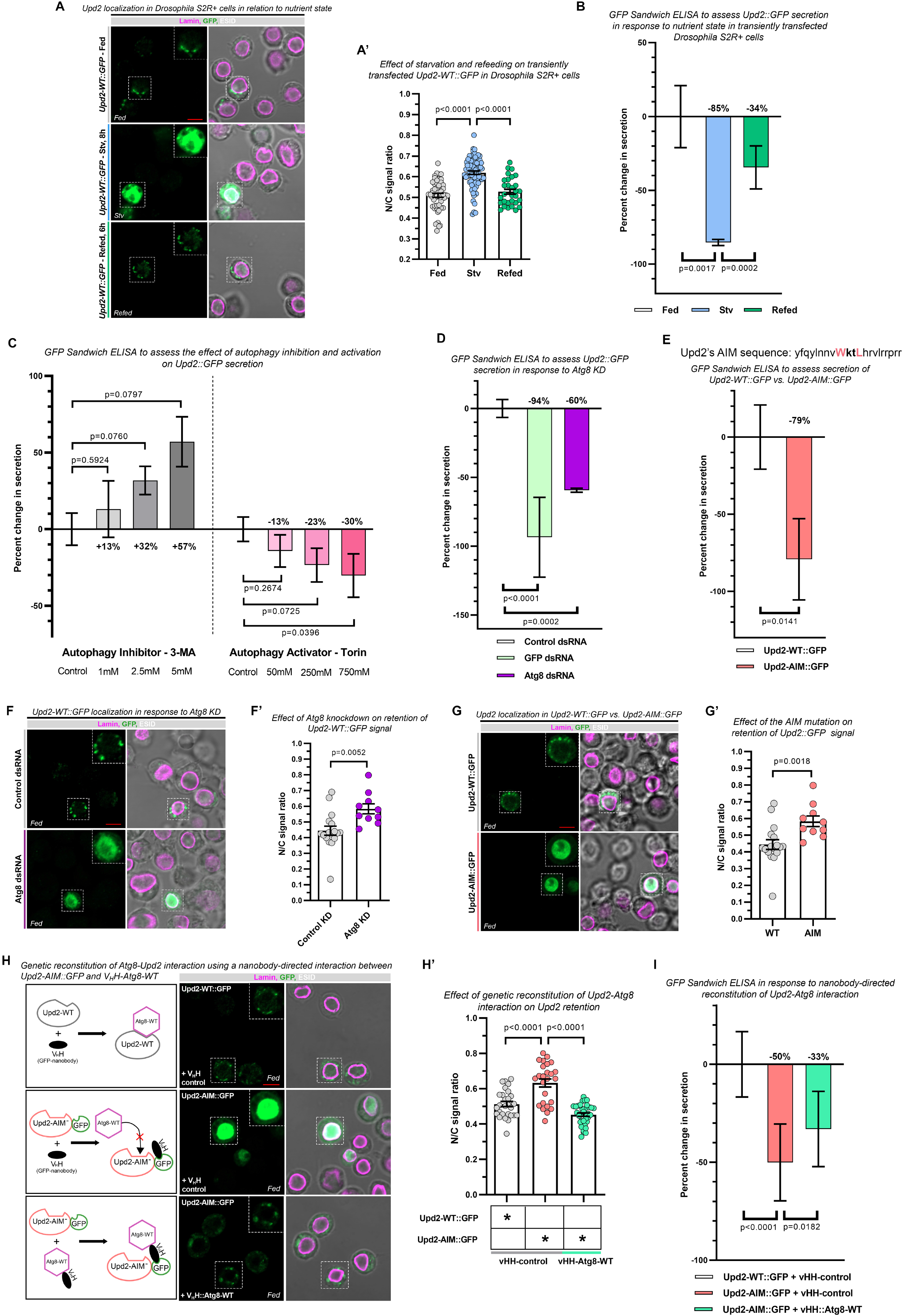
Upd2’s interaction with Atg8 governs its secretion in *Drosophila* S2R+ cells. A) Confocal micrographs of single optical slices of *Drosophila* S2R+ cells transiently transfected with Upd2-WT::GFP (green; anti-GFP) and co-stained with Lamin (magenta) in complete media (upper panel), 8hrs nutrient-deprived (middle panel) and refed with complete media (lower panel). In A’, the ratio of Upd2::GFP nuclear/whole cell intensity is plotted. Data is shown as mean ± SEM. Each dot represents a cell, 30-80 cells were counted per treatment condition. Statistical significance was measured by a non-parametric two-tailed Mann Whitney U test. B-E) A GFP sandwich ELISA assay was performed on conditioned media of S2R+ cells that were transiently transfected with Upd2-WT::GFP to quantitatively assess Upd2 secretion. Cells were exposed to differing nutrient conditions (B), drugs (C), dsRNA (D), or an Upd2 mutant (E). Normalized percent fold change of the secreted GFP signal is plotted. Error bars represent %SD or %SEM. Statistical significance was measured by an unpaired two-tailed t-test on 6 biological replicates per condition. F-H) Confocal micrography of single optical slices was performed for *Drosophila* S2R+ cells transiently transfected with Upd2-WT::GFP (green), stained with Lamin (magenta), and co-transfected with either indicated dsRNA (F), Upd2 point mutations (G) or co-transfected vHH nanobody-tagged constructs; note that a schematic of experimental design is shown on the left column (H). Scale bar is 5um. Each dot in the corresponding quantitation (F’, G’, H’) represents a cell. 10-40 cells were counted per condition. Data is shown as mean ± SEM. Statistical significance was measured by an unpaired two-tailed t-test or a non-parametric two-tailed Mann Whitney U test. I) A GFP sandwich ELISA assay was performed on conditioned media of S2R+ cells that were transiently co-transfected with Upd2-WT::GFP+V_H_H, Upd2-AIM::GFP+V_H_H or Upd2-AIM::GFP+V_H_H-Atg8-WT to quantitatively assess Upd2 secretion. Normalized percent fold change of the secreted GFP signal is plotted. Error bars represent %SD. Statistical significance was measured by an unpaired two-tailed t-test on 6 biological replicates per condition.

To determine if Upd2’s starvation-induced intracellular accumulation correlates with reduced secretion, we performed a quantitative ELISA for GFP-tagged Upd2^37^. In a prior study^38^, we identified conserved factors required for Upd2 and Leptin secretion in Drosophila S2R+ cells using this assay. We observed that 8 hours of AA deprivation caused a significant reduction in Upd2 secretion (85%; p=0.0017), and refeeding for 6 hours significantly ameliorated this effect (p=0.0002) (**Fig. 1B**). These results suggest that *Drosophila* cells possess cell-intrinsic mechanisms that couple Upd2 secretion to nutrient state.

A well-documented cellular response to starvation is the upregulation of the nutrient scavenging pathway, Autophagy ^39^. We wondered whether autophagy induction might play a role in restricting adipokine secretion as the two processes represent responses to opposite nutrient states. On pharmacological inhibition of autophagy, using 3-Methyladenine (3-MA), we observed a dose-sensitive increase in Upd2 secretion (**Fig. 1C**; 5mM resulted in a 57% increase; p=0.08). Conversely, treatment with Torin1, which activates autophagy^40^, reduced Upd2 secretion (**Fig. 1C**; 750nM causes 30% reduction; p=0.04). In *Drosophila* S2R+ cells, AA deprivation potently induces nutrient deprivation signals^35,36^. These results suggest that autophagy induction is antagonistic to Upd2 secretion.

LC3, a key player in autophagy, has also been implicated in LC3-dependent extracellular vesicle loading and secretion (LDELS)^27^. In the LDELS pathway, LC3 and its associated components play a role distinct from their established roles in classical autophagy. Given that autophagy was found to be antagonistic to Upd2 secretion (**Fig. 1C**), we hypothesized that LC3/Atg8 controls Upd2 secretion from well-fed cells. In support of our hypothesis, we observed that dsRNA-mediated knockdown of Atg8 results in a significant reduction in Upd2 secretion (p=0.0002) (**Fig. 1D**), as measured by a quantitative GFP sandwich ELISA.

During Autophagy, Atg8 recruits specific proteins to the autophagosome via a short linear protein domain termed the LC3/Atg8-interacting motif (LIR/AIM). This motif contains the consensus sequence W/F/Y-X-X-L/I/V^41^. Recently, the AIM/LIR sequences have been shown to be required for LC3 loading of cargos-such as RNA-binding proteins (RBPs) - into EVs during LDELS-based secretion^27^. Point mutations to the RBP’s LIR sequence are sufficient to disrupt RBP’s LDELS-mediated secretion^27^. Hence, we asked if *Drosophila* Upd2, like RBPs, might represent direct cargoes for Atg8. We analyzed protein sequences and found that Upd2 (**Fig. 1E**) has a putative AIM sequence. We found that single amino acid point mutations of the AIM motif of Upd2 (W and L to A) reduced secretion by 79% (p=0.01) (**Fig. 1E**).

Examining the intracellular localization of Upd2 in Atg8-KD (**Fig. 1F, F’**) and Upd2-AIM (**Fig. 1G, G’**), we observed increased Upd2 intracellular accumulation. These observations were consistent with the AA starvation experiments in which we observed reduced secretion (Fig. 1B) and increased Upd2 intracellular accumulation (**Fig. 1A, A’**). These results suggest that Atg8 promotes Upd2 secretion via interaction between Atg8 and Upd2’s AIM sequence.

We reasoned that the increased intracellular accumulation and decreased secretion of Upd2-AIM were due to its inability to interact with Atg8. If true, restoring its interaction with Atg8 would be sufficient to rescue the intracellular accumulation phenotype and enhance the secretion of Upd2-AIM. To address this, we employed a nanobody-based approach (vhhGFP4 – nanobody to GFP) to engineer interactions between Upd2-AIM and Atg8 (See Schematic in **Fig. 1H**)^42^. This approach has been used to investigate how the interaction between two proteins, one tagged with V_H_H and the other with GFP, alters localization and protein behavior^43-45^. Therefore, we tagged Atg8 with V_H_H (V_H_H::Myc-Atg8-WT) and co-expressed it with Upd2-WT::GFP and Upd2-AIM::GFP. We observed that co-expression of the control (V_H_H::Myc control) does not result in localization change of Upd2-WT (**Fig. 1H, top panel**), nor does it reduce the intracellular accumulation of Upd2-AIM (**Fig. 1H, middle panel**). But in the presence of V_H_H-tagged Atg8-WT (V_H_H::Myc-Atg8-WT), we observed that Upd2-AIM accumulation was significantly reduced and localized to cytosolic puncta akin to Upd2-WT (**Fig. 1H bottom panel, H’**). This suggests that Upd2-AIM’s interaction with Atg8 is sufficient to restore its intracellular accumulation to wild-type levels.

Next, we examined the secretory potential of Upd2-AIM reconstituted with Atg8. As expected, Upd2-AIM secretion was significantly compromised compared to Upd2-WT when co-expressed with the V_H_H::Myc control (**Fig. 1I**; 50% reduced; p<0.0001). Reconstitution with Atg8 significantly improved the secretory capacity of Upd2-AIM relative to control **(Fig. 1I**; p=0.0182). While Upd2-AIM’s secretion potential improves on its reconstitution with Atg8 compared to the control nanobody (**Fig. 1I**), it is not restored to WT levels. Overall, nanobody-based reconstitution between Atg8 and Upd2-AIM shows rescue of Upd2-AIM intracellular accumulation defects. Hence, our results are consistent with the interpretation that Upd2-AIM’s defective interaction with Atg8 underlies its intracellular retention in *Drosophila* S2R+ cells.

### Atg8 controls Upd2 intracellular accumulation in adult *Drosophila* adipocytes

To validate our *in vitro* cell biological findings *in vivo*, we generated transgenic flies with a fat-tissue-specific expression of GFP-tagged Upd2-WT (*Lpp-Gal4>UAS-Upd2-WT::GFP*) and Upd2-AIM (*Lpp-Gal4>UAS-Upd2-AIM::GFP*) and performed image-based quantitative analysis of Upd2 accumulation in fly fat. Upd2-WT::GFP is detected at very low levels in adult Drosophila adipocytes despite over-expression. In contrast, over-expression of *Upd2-AIM* in adult fly fat is readily detectable, and displays increased intracellular localization (**Fig. 2A, A’**; p=0.0400). Strikingly, co-expression of V_H_H-tagged Atg8 with Upd2-AIM in the adult fly fat reduces Upd2-AIM intracellular accumulation to Upd2-WT levels (**Fig. 2A-bottom panel, A’**; p=0.0061). Hence, in adult *Drosophila* abdominal adipocytes, Upd2-AIM mutation leads to its intracellular retention, and nanobody-based reconstitution with Atg8 reduces the intracellular accumulation of Upd2-AIM.

**Figure 2.**
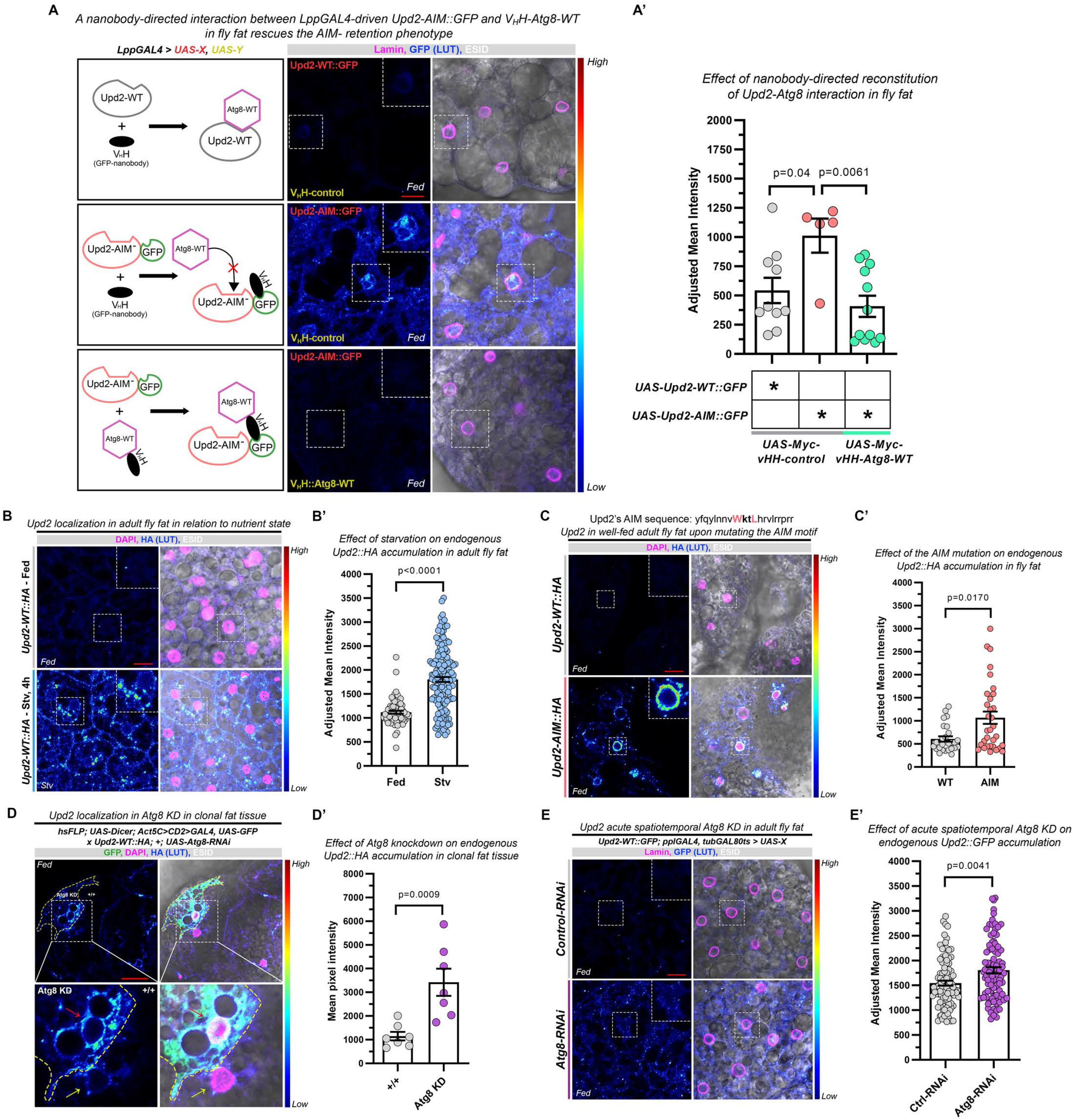
*In vivo Drosophila* model to study Atg8’s role in regulating Upd2’s intracellular retention. Confocal micrography of single optical slices was performed for adult *Drosophila* abdominal fat tissue. Samples were dissected, fixed and immunostained with antibodies against GFP (rainbow-LUT) (A), HA (rainbow-LUT) (B-E), Lamin (magenta) (A, E) or DAPI (magenta) (B-D). Scale bar is 10um for A-C, E; and 20um for D. Each dot in the corresponding quantitation (A’-E’) represents either a cell (A’-C’, E’) or an area demarcated by a GFP filter (D’). Data is shown as mean ± SEM. Statistical significance was measured by an unpaired two-tailed t-test or a non-parametric two-tailed Mann Whitney U test. A) Transgenic flies co-expressing UAS-Upd2::GFP (WT or AIM) along with UAS-VHH (control or Atg8-WT) under the control of a fat-specific LppGAL4 driver were generated. A schematic describing the experiment is shown in the first column. Adjusted mean intensity of the GFP-LUT signal from 5-12 cells was quantified per condition in A’. B) Upd2 genomic locus is tagged with HA endogenously using CRISPR technology (referred to as Upd2-WT::HA). Abdominal fat from well-fed 7-day old male flies (top panel) and 4 hours acute starvation (bottom panel). Mean intensity is quantified in B’. 71-158 cells were quantified per condition. C) CRISPR technology was used to generate flies in which the endogenous Upd2’s AIM motif was mutated from WKTL to AKTA (referred to as Upd2-AIM::HA). Abdominal fat from age matched male flies WT (top panel) and AIM (bottom panel), quantified in C’. 26-32 cells were quantified per condition. D) FLP-OUT technique was used to generate Act-GAL4-driven GFP-positive clones in adult fly fat in the background of the Upd2-WT::HA CRISPR knock-in. The top panel shows the effect of Atg8 knockdown on HA-LUT signal in a positive clone (demarcated by a yellow dashed line), and bottom panel shows a higher magnification of the inset. The Upd2-HA-LUT signal within the positive knockdown of the clone relative to the surrounding wild-type tissue is quantified in D’. 7 clones were quantified to assess their significance. The corresponding control knockdown is shown in Supplemental Figure S2. E) Acute spatiotemporal knockdown of Atg8 in adult fly fat (bottom panel) compared to control-RNAi knockdown (top panel) and quantified in E’. Signal from 84-97 cells was quantified per condition.

To investigate the effect of acute starvation on Upd2 endogenous retention *in vivo*, we used CRISPR-Cas9 technology to generate a tagged knock-in, such that the endogenous Upd2 gene was tagged with HA at the N-terminus. We subjected these *Upd2-WT::HA* flies to 4 hours of acute starvation. Using anti-HA immunostaining, we found that Upd2 is almost undetectable by confocal imaging at baseline fed state (**Fig. 2B**). However, upon starvation, we observed a significant increase in Upd2 levels within the adult abdominal adipose tissue. (**Fig. 2B, B’**; p<0.0001). Hence, similar to its vitro regulation, Upd2 accumulates within adult fly adipocytes during acute starvation.

We also generated knock-in point mutations in Upd2’s AIM sequence in the endogenous HA-tagged Upd2 locus. The homozygous *Upd2-AIM::HA* flies were viable and fertile. We examined the localization of the mutant adipokine, Upd2-AIM, via confocal microscopy and found a significant increase in its intracellular accumulation (**Fig. 2C, C’**; p=0.0170).

Next, we tested whether the genetic knockdown of Atg8 phenocopies the Upd2-AIM point mutations. We generated genetic mosaic tissue in *Drosophila* adipocytes using the FLP-out Gal4 technique^46^. Using this tool, we expressed *Atg8-RNAi* in a small subset of *Drosophila* adipocytes (GFP-positive) surrounded by wild-type tissue expressing normal, endogenous levels of Atg8 (GFP-negative). We observed a significant accumulation of HA-tagged endogenous Upd2 only in patches of tissue where Atg8 was knocked-down (**Fig. 2D, D’**; p=0.0009) (note GFP-positive cells express Atg8-RNAi). Importantly, the expression of RNAi alone (control-*Luciferase-RNAi)* in a subset of clonal adipose tissue did not result in increased Upd2 accumulation (**Fig. S2**; p=0.3699). Hence, the knockdown of Atg8 in a subset of cells within a fat tissue is sufficient to increase Upd2 retention. In a second line of experiments, we acutely knocked down Atg8 in the adult fat using the tubGal80ts^47^ system for a week and assayed Upd2 endogenous accumulation. We observed increased Upd2 accumulation when Atg8 is knocked down compared to control RNAi (**Fig. 2E, E’**; p=0.0041). Altogether, our experiments establish that Atg8 knockdown, and points mutations to Upd2’s AIM, increase Upd2 retention in well-fed adipocytes.

### Upd2 intracellular accumulation promotes organismal starvation resilience

In mammals and flies, reduced adipokine leptin/Upd2 during starvation allows the neuroendocrine system to adapt to a nutrient-deprived state^1,7,48 2^. Since *Upd2-AIM* mutation increases its intracellular accumulation, independent of nutrient state, it presents a genetic model to test the effects of constitutive adipokine retention on survival and the physiological adaptation to starvation.

To this end, we performed a fat-tissue-specific over-expression of the Upd2-AIM transgene in a *Upd2Δ* mutant background (*Upd2Δ; Lpp-Gal4> UAS-Upd2-AIM)* and compared it with overexpression of a control empty transgene (*UAS-Luciferase*) and Upd2-wildtype transgene (*UAS-Upd2-WT*). We found that flies expressing control transgene live longer on a 1% sucrose-agar-starvation diet than Upd2-WT over-expressing flies (**Fig. 3A**; p=0.0216). This is consistent with our previous reports that *Upd2Δ* is starvation-resistant^2^. Strikingly, we found that adipocyte-specific over-expression of Upd2-AIM in a *Upd2Δ* background significantly increased survival, even more than the *Upd2Δ* loss-of-function allele (**Fig. 3A**; p<0.0001). This suggests that AIM-mediated retention improves upon the starvation resilience of the loss-of-function allele. Since we utilized fat-tissue-specific overexpression, we were curious to determine whether a similar effect might be observed when the AIM mutation was expressed at an endogenous level. Hence, we compared the survival capacity of CRISPR-engineered AIM mutation at the endogenous Upd2 locus (*Upd2-AIM::HA*) with a wild-type control (*Upd2-WT::HA)* on prolonged starvation. As with the overexpression data, we observed an improvement in survival with the CRISPR-generated AIM mutant relative to the WT control (**Fig. 3B**; p=0.0005).

**Figure 3.**
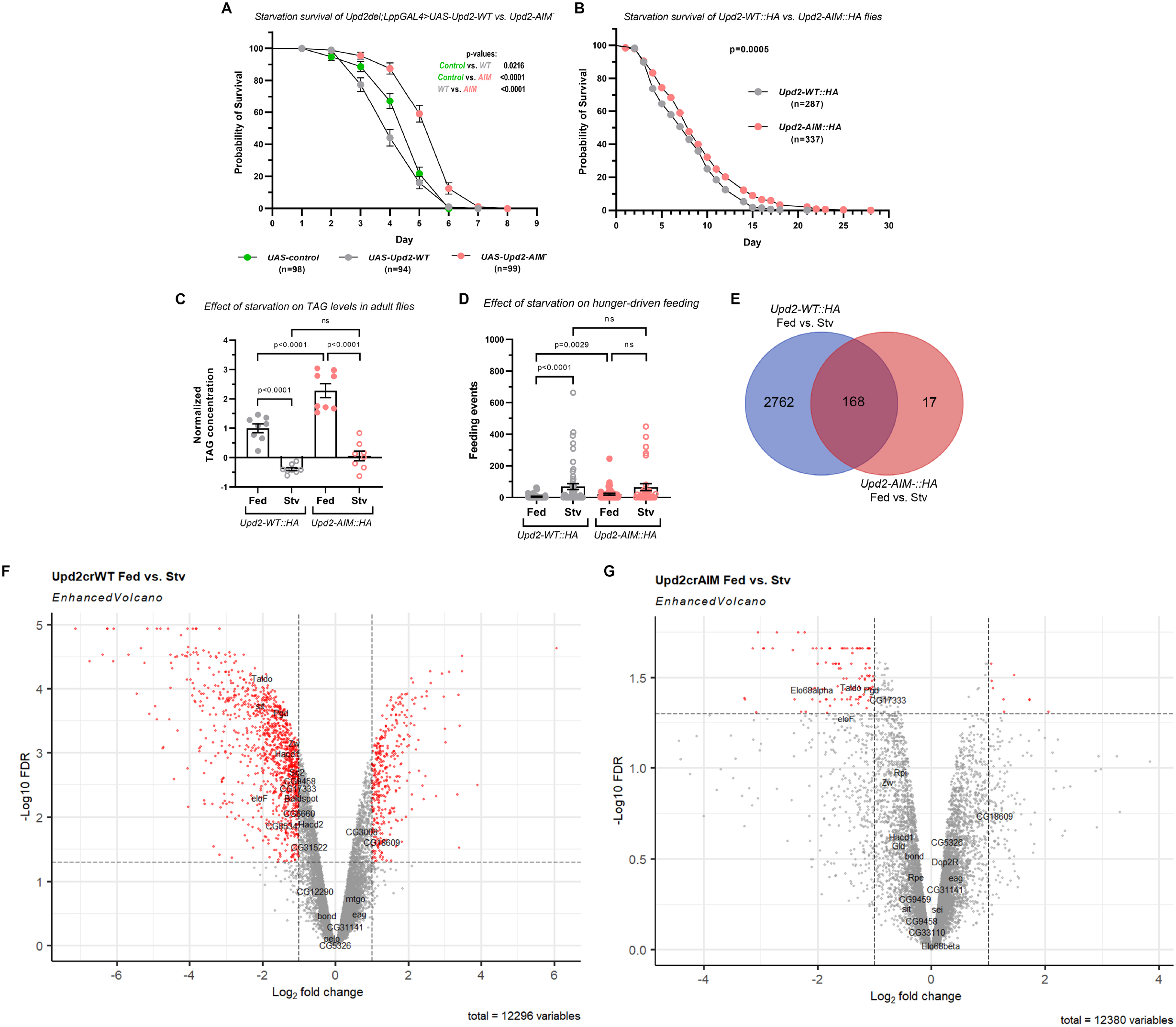
Upd2’s intracellular retention confers a survival advantage upon starvation, due to large scale transcriptomic alterations. A, B) Survival curves of indicated genotypes on 1% sucrose agar starvation media. In A) fat-specific LppGAL4 driver was used to over-express a control transgene (grey), Upd2-WT (green), or Upd2-AIM (salmon) in an Upd2-deletion background. In B) starvation survival of endogenously HA tagged Upd2-WT::HA (grey) and Upd2-AIM::HA (salmon) flies was compared. C) Triacylglycerol (TAG) levels of flies were quantified for Upd2-WT::HA vs Upd2-AIM::HA CRISPR knock-in flies in fed states (ad libitum standard diet) compared to starvation states (16 hours; 0% sucrose). Each dot represents one sample (three whole adult male flies/sample) with n=8. D) Feeding events were measured for Upd2-WT::HA vs Upd2-AIM::HA CRISPR knock-in flies in fed states (ad libitum standard diet) versus starvation states (16 hours; 0% sucrose) using the FLIC assay. Each dot represents the feeding events of an individual fly (n=32-59). Following a failed normality test, a Kruskal-Wallis test with Dunn’s multiple comparisons was performed post-hoc. E) Venn diagram depicting the number of differentially expressed genes in Fed vs. Stv state for Upd2-WT::HA flies (blue circle), compared to the number of differentially expressed genes in Fed vs. Stv state for Upd2-AIM::HA flies (red circle). The number of genes whose transcription is altered as a function of nutrient state in Upd2-WT::HA flies (n=2930) compared to Upd2-AIM::HA flies (n=185) is indicated. F, G) RNASEQ analysis was conducted on four samples (Upd2-WT::HA Fed, Upd2-WT::HA Stv, Upd2-AIM::HA Fed, and Upd2-AIM::HA Stv) to compare changes in global gene expression as a function of different nutrient conditions and genotypes. Volcano plot showing differential gene expression in Fed vs. Stv Upd2-WT::HA flies (F) and Upd2-AIM::HA (G); red Dots indicate significantly differentially expressed genes (FDR <0.05) with a log fold change greater than +/-1. See Figure S3 for GO plots.

To ascertain a reason for this extended survival, we analyzed the neutral fat stores of *Upd2-AIM::HA* flies by measuring whole-body triglyceride (TAG) levels. *Upd2-AIM::HA* flies displayed significantly higher TAG levels than the wild-type controls (**Fig. 3C**; p<0.0001). However, when subjected to overnight starvation, they efficiently lipolysis their fat stores (**Fig. 3C**; p<0.0001). This suggests that constitutive accumulation of Upd2 within *Drosophila* fat, unlike a *Upd2Δ* loss of function allele, extends resilience at least in part by storing excess energy reserves as neutral fat and then efficiently breaking it down during starvation. Thus, *Upd2-AIM::HA* behaves like a neomorphic gain-of-function mutation, as its starvation behavior and fat physiology differ from the loss-of-function *Upd2Δ*^2^.

To understand why *Upd2-AIM::HA* flies had higher baseline TAG reserves, we assessed feeding behavior using the FLIC (Fly Liquid-Food Interaction Counter) assay^49^. We observed an increased number of feeding events in *Upd2-AIM::HA* relative to wild-type controls (**Fig. 3D**; p=0.0029), which could be a possible explanation for their elevated TAG reserves (**Fig. 3C**). As a response to starvation, animals exhibit a huger-driven feeding response. As expected, we observed a significant increase in feeding events in *Upd2-WT::HA* flies after they were starved overnight (**Fig. 3D**; p<0.0001). Interestingly, there was no difference in feeding behavior upon starvation in *Upd2-AIM::HA* flies (**Fig. 3D**). Hence, the feeding behavior of *Upd2-AIM* flies at baseline mimics the starved state.

The observation that Upd2 retention within adipocytes increases starvation resilience suggests that it might rewire systemic metabolism toward a state better adapted to starvation. To understand what these pathways are likely to be, we surveyed the transcriptomes of *Upd2-AIM::HA* and *Upd2-WT::HA* flies by performing bulk RNAseq. We analyzed both genotypes’ differentially expressed (D.E.) genes between fed and starved states. In *Upd2-WT::HA* flies, 2930 genes were altered during overnight starvation (**Fig. 3E, F**). However, it was striking that the *Upd2-AIM::HA* flies showed only 185 DE genes (7% of the changes in WT state; **Fig. 3E, G**) upon starvation. An interpretation of the minimal changes we observe in the transcriptome of flies in which Upd2 is constitutively retained, despite its nutrition state, is that it mimics a biological state that prewires it to starvation. This transcriptomics analysis is consistent with the survival analysis shown in **Fig. 3B**, which experimentally demonstrates the starvation resilience of *Upd2-AIM* flies.

We performed Gene Ontology (GO) analysis^50,51^, followed by GO enrichment^52^, to identify pathways that were differentially regulated between the genotypes. In control (*Upd2-WT::HA*) flies, pathways involved in reproductive physiology, innate immunity, and cellular metabolism (specifically fatty acid biosynthesis and pentose phosphate shunt) were altered (**Fig. S3A**), whereas in *Upd2-AIM::HA* flies, the pentose phosphate pathway was most significantly altered between fed and starved states (**Fig. S3B**). The transcriptomics survey leads us to propose that Upd2 retention rewires a metabolic shift towards adaptations that allow for better starvation survival.

### LC3 family proteins promote human leptin secretion

Human leptin can functionally substitute for Upd2 function in flies^2^. The high degree of evolutionary conservation between *Drosophila* Atg8 and mammalian LC3-family proteins and the autophagy pathway are well-established^53-56^. Given this, we explored whether LC3-GABARAP regulates leptin secretion. Human leptin has two putative AIM/LIR sequences near its N-terminus (Fig. 4A). We generated point mutations to these two motifs (AIM1: WgtL→ AgtA; AIM2: WpyL→ ApyA) and transiently transfected Leptin-WT::GFP and Leptin-AIM::GFP plasmids into HEK293T cells. Since HEK293T cells do not express endogenous leptin, we could compare the effect of the AIM point mutations on leptin secretion in the conditioned media of HEK293T cells using a quantitative GFP sandwich ELISA. We found that compared to baseline Leptin-WT::GFP, the AIM mutations caused a reduction in secretion for AIM1 (50% reduction; p= 0.1367) and a statistically significant reduction in secretion for AIM2 (72% reduction; p=0.0228). On mutating both AIM sequences, we observed a significant (89% reduction; p=0.0039) in leptin secretion (**Fig. 4A**). This suggests that leptin’s putative AIM sequences, like in Upd2, are important for secretion.

**Figure 4.**
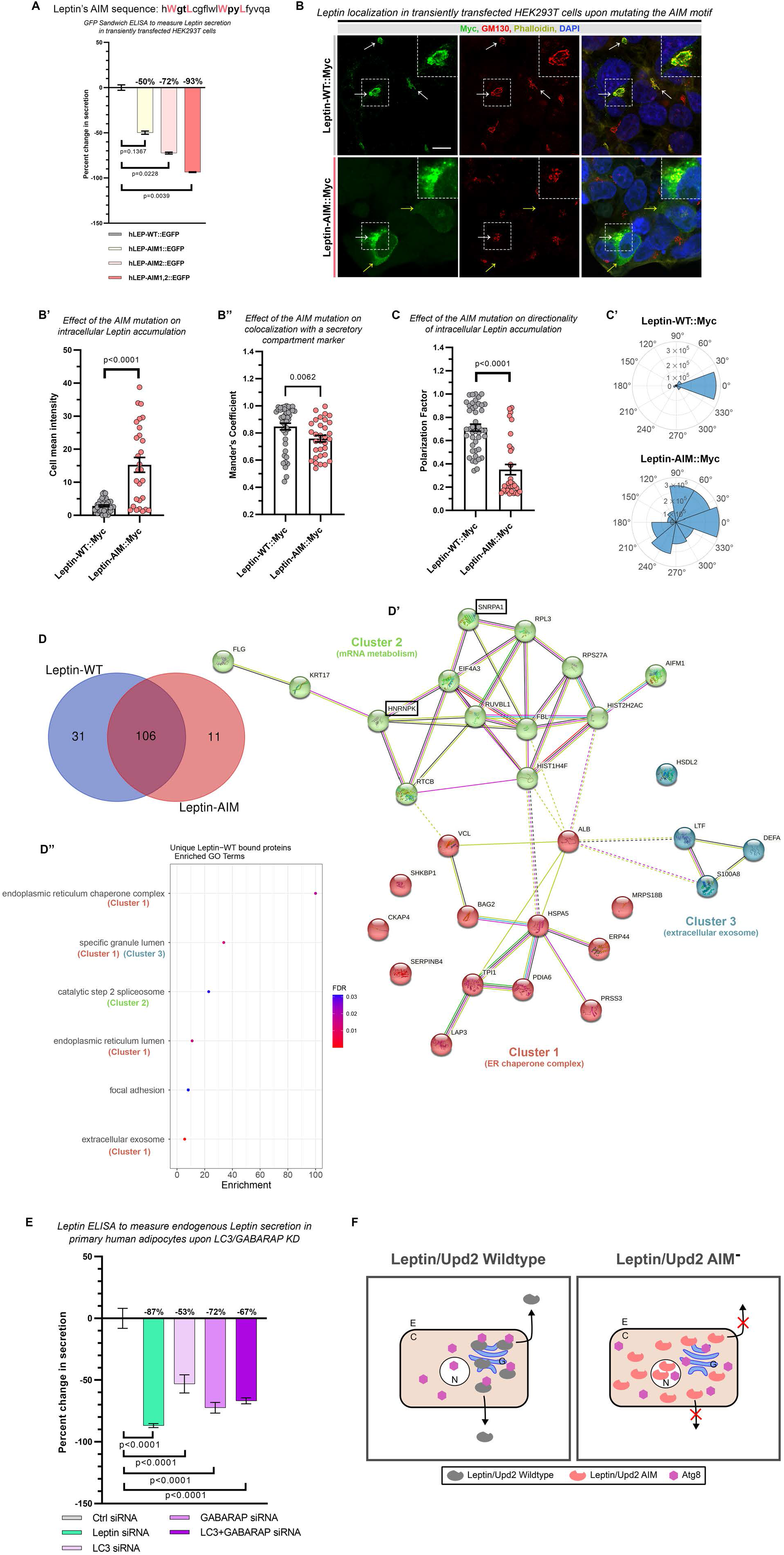
LC3/GABARAP is required for Leptin secretion in human cells. A) A GFP sandwich ELISA assay was performed on conditioned media of HEK293T cells that were transiently transfected with hLep-WT::EGFP, hLep-AIM1::EGFP (WGTL→AGTA), hLep-AIM2::EGFP (WPYL→ APYA) to quantitatively assess Leptin secretion. Normalized percent fold change of the secreted GFP signal is plotted, with hLep-WT::EGFP (grey) as the control. Error bars represent %SD. Statistical significance was measured by an unpaired two-tailed t-test on 6 biological replicates per condition. B) Confocal micrography of single optical slices was performed for HEK293T cells, that were transfected with a Myc-tagged Leptin-WT (top panel) or Leptin-AIM (both AIMs mutated; bottom panel). Cells were fixed and immunostained with anti-Myc (green), anti-GM130 (red) and DAPI (blue). Leptin’s intracellular accumulation (quantified in B’), and colocalization with GM130 (quantified in B’’). Scale bar is 10um. Each dot in the corresponding quantitation represents a cell. 28-44 cells were counted per condition. Data is shown as mean ± SEM. Statistical significance was measured by a non-parametric two-tailed Mann Whitney U test. C) A polarization histogram for Leptin::Myc was calculated to describe and quantify its distribution and is pictorially represented in C’. Each dot represents a cell. 32-44 cells were counted per condition. Data is shown as mean ± SEM. Statistical significance was measured by a non-parametric two-tailed Mann Whitney U test. Also see Supplemental Figure 4A for additional plots. D) A Venn diagram depicting the number of interacting partners identified in an IP-MS screen of Leptin-WT::GFP vs Leptin-AIM::GFP. HEK293T cells were transfected with either Leptin-WT::GFP or Leptin-AIM::GFP, and a GFP pulldown followed by a mass spectrometric analysis was done to investigate differences in Leptin’s interactome. D’ shows STRING analysis of the 31 proteins that were unique interactors of Lepin-WT. D’’ lists the GO terms of proteins associated with Leptin-WT. Enrichment for the GO term is displayed on the X-axis and the color of the dot represents the significance, with blue indicating an FDR=0.05 and red indicating an FDR<0.0001. E) Endogenous Leptin ELISA from primary human adipocytes that incubated with either nontargeted siRNA (positive control; grey), Leptin siRNA (negative control, green), LC3 siRNA (mauve), GABARAP siRNA (lilac), and LC3 + GABARAP siRNA (purple). siRNA treatment did not affect adiposity of the cells (See Supplemental Figure 4). Error bars represent %SD. Statistical significance was measured by an unpaired two-tailed t-test on 6 biological replicates per condition. F) Summary model. Leptin/Upd2 secretion requires LC3/GABARAP: A conserved model for adipokine secretion from human adipocytes to fly fat, showing that Leptin/Upd2 (adipokine) requires interaction with LC3/GABARAP/Atg8 (transporter) to be efficiently secreted from adipocytes to signal nutrient status to the brain. Disrupting the interaction of the adipokine with its transporter results in intracellular retention, reduced secretion, and impaired signaling to the brain resulting in large scale transcriptomic/metabolic alterations.

Next, we examined the localization of Leptin-AIM::GFP versus Leptin-WT::GFP control in HEK293T cells using imaging-based approaches. We found that Leptin-WT::GFP localizes asymmetrically within the cell to a perinuclear region that is positive for the secretory compartment marker GM130^57^ (**Fig. 4B**, see polarization plots in **4C’**; **Fig. S4A**). In contrast, mutations to both of leptin’s AIM motifs result in increased leptin accumulation (**Fig. 4B’**; p<0.0001), reduced co-localization with GM130 (**Fig. 4B"**; p=0.0062), loss of directionality of leptin accumulation (**Fig. 4C, C’**; p<0.0001; **Fig. S4A, A’**), and an overall diffuse cytosolic localization (**Fig. 4B**). Our observations are consistent with the interpretation that mutations to leptin’s AIM sequences interfere with its ability to be directed to an appropriate secretory compartment.

We performed comparative analyses of protein-protein interactors between WT and AIM leptin to investigate how leptin is secreted. We immunoprecipitated (IP) Leptin-WT::GFP and Leptin-AIM::GFP from HEK293T cells and subjected the immunocomplexes to LC-MS. We then performed unbiased proteomic analyses to identify proteins present specifically in the secretion-competent leptin-WT and absent in the leptin-AIM pulldown, which we further analyzed using the STRING database^58,59^. Based on our analysis cutoff, we found that 31 proteins bound uniquely to leptin-WT but not to leptin-AIM (**Fig. 4D**). These proteins fell into three functional interactions clusters (**Fig. 4D’, D"**) – Cluster 1 (E.R. chaperone complex), Cluster 2 (mRNA metabolism) and Cluster 3 (extracellular exosome). G.O. enrichment analysis showed that the following G.O. terms displayed a 10-100-fold enrichment only with leptin-WT: endoplasmic reticulum (E.R.) chaperone, granule lumen, extracellular exosome, RNA binding proteins (RBPs-spliceosomes) (**Fig. 4D"**). We were intrigued to observe that 13 proteins involved with mRNA metabolism were enriched only in leptin-WT immunocomplexes (**Fig. 4D’**). Specifically, we noted that secretion-competent leptin associates with two RNA binding proteins (**Fig.- 4D’**-green) — HNRNPK and SNRPA1. These RNA-binding proteins are secreted by LC3-dependent extracellular-vesicle loading and secretion (LDELS)^27^. Hence, we postulated that LC3 regulates human leptin secretion, probably via an LDELS-like route.

We further examined the requirement for LC3 family proteins in endogenous leptin secretion from human primary subcutaneous adipocytes to test this in a physiologically relevant system. We performed knockdown of endogenous LC3s (LC3A, B, C) and GABARAPs (GABARAP, GABRAPL1, GABRAPL2) using siRNAs that were validated in previous studies^60^, and performed an ELISA for endogenous leptin. As a positive control for a knockdown, we used leptin siRNA. We observed a statistically significant reduction in endogenous leptin secretion from human subcutaneous adipocytes upon knockdown of LC3 (p<0.0001), GABARAP (p<0.0001), or both (p<0.0001) (**Fig. 4E, S4B**). The treatment of adipocytes with siRNAs did not affect the adiposity of the cells, which was measured via AdipoRed analysis (**Fig. S4B’**). Hence, we propose that Atg8/LC3 & GABARAP proteins play an evolutionarily conserved role in promoting the secretion of Upd2 and leptin from *Drosophila* and human adipocytes (**Fig. 4F**).

## Discussion

We establish here a novel evolutionarily conserved role for Atg8/LC3 proteins in promoting Upd2/Leptin secretion across multiple levels of biological complexity – *in vitro* insect and mammalian cell systems, *in vivo* fly adipocytes, and cultured human adipocytes. It is well-recognized that Atg8 and LC3 proteins enable organismal adaptation to starvation because of their conserved roles in autophagy from yeast to mammals^61-63^. On starvation, Atg8 is recruited to autophagosomes^18^, and its association with double membranes and subsequent fusion steps with the lysosome is likely to limit its availability for leptin secretion. Hence, due to Atg8/LC3’s involvement in autophagy-related functions, Upd2/leptin would be retained within cells on starvation. In support of this model, we observed that activation of autophagy inhibits adipokine Upd2 secretion (**Fig. 1C**). The role of LC3 protein in nutrient deprivation has been long appreciated^64^. Now, our findings establish Atg8/LC3-mediated adipokine secretion control as a cell-autonomous mechanism regulating organism-wide responses to nutritional flux.

A key technical challenge of studying the adipokine, Upd2, in *Drosophila* has been the lack of robust means to quantify Upd2 secretion in the fly hemolymph. Even sophisticated mass spectrometry analysis cannot detect Upd2 in fly hemolymph^65^. Hence, we adopted a strategy analogous to studies on *Drosophila* insulin-like peptides (Dilps). In these studies, retention of Dilp2 and Dilp5 within insulin-producing cells (IPCs) indicates reduced secretion^66,67^. In addition, we have performed quantitative secretory assays on *Drosophila* S2R+ cells to investigate Upd2 secretion. Indeed, a tight inverse correlation between secretion and intracellular Upd2 retention is observed *in vitro* (**Fig. 1A, B**). Hence, for *in vivo* studies, as published in previous work from our lab^38^, we use Upd2 accumulation within adult *Drosophila* adipocytes as a proxy for secretion.

In *Drosophila* S2R+ cells AA-deprivation potently induces nutrient deprivation signals^35,36^. Consistent with this, we find AA deprivation causes Upd2 retention and reduced secretion, but culturing S2R+ cells in serum-free media does not have this effect. We postulate that this is due to a tight link between autophagy induction and AA deprivation. Application of the autophagy activator, Torin1, significantly reduces adipokine Upd2 secretion (**Fig. 1C**), providing further evidence to support that autophagy activation is antagonistic to adipokine secretion. Altogether, our study suggests an intriguing possibility that Atg8/LC3’s bidirectional role in autophagy and secretion is likely to exquisitely fine-tune Upd2/leptin release to communicate cell-intrinsic nutrient state.

*Upd2-AIM* behaves like a neomorphic gain-of-function mutation, as its feeding behavior and fat physiology differ from the loss-of-function *Upd2Δ^2^*. *Upd2-AIM* flies have elevated baseline TAG levels and display increased baseline feeding behavior compared to *Upd2-WT*. A recent study that artificially tethered Upd2 to adult adipocyte membranes to prevent its secretion found that baseline TAG levels were much higher in such flies^68^. Hence, it appears that constitutive Upd2 retention increases baseline TAG levels, at least in part via elevated feeding behavior in baseline. Constitutive adipokine Upd2 retention results in physiological changes in *Upd2-AIM* flies-increased baseline feeding, elevated TAG stores and a transcriptome signature that is closer to starved state-these changes are likely to allow *Upd2-AIM* flies display increased starvation resilience.

The transcriptome survey of *Upd2-AIM* on starvation suggests that constitutive Upd2 retention acts as a "starvation-like" signal to rewire metabolism to better adapt to starvation. In *Upd2-AIM* flies, there are much fewer transcriptome changes (∼180-7% of changes observed in WT) on starvation compared to >2700 genes that change in wild-type flies. On examining the baseline expression of PPP genes in Upd2-AIM flies compared to wild-type flies, we observed that a few PPP genes (*CG7140, Rpe)* were already downregulated even in the fed state (**Fig. S3C**). Recent studies in yeast have shown that a novel adaptation strategy to prolonged glucose deprivation is to downregulate the pentose phosphate pathway (PPP); this allows energy can be conserved instead of generating NADPH for nucleotide precursors^69^.

Prior studies have examined how leptin secretion and production are controlled. In rat adipocytes, insulin stimulates post-prandial leptin secretion^70^, and transcription^71,72^. The mTOR pathway positively regulates leptin mRNA translation in 3T3-L1 adipocytes^73^. Furthermore, in human abdominal and omental fat deposits, TNF and glucocorticoids increase leptin expression during obesity and stress^74^. Studies in 3T3-L1 and rat primary adipocytes suggest leptin is enriched in the endoplasmic reticulum and traffics to the Golgi^75-77^. However, independent studies report leptin does not traffic via known adipocyte secretory routes such as GLUT4-positive vesicles^75-77^. In rat adipocytes, leptin is present in low-density small intracellular vesicular compartments^78^. But, the nature of the leptin-positive "light" secretory vesicles, and the cell-intrinsic factors that direct leptin to the appropriate secretory compartment, are yet to be elucidated. Based on our unbiased proteomic analyses, we propose that leptin is secreted by LC3-mediated exosome secretion. In the future, it will be important to tease apart whether LC3-mediated leptin secretion involves recently established forms of Atg8 lipid conjugation to single membranes-such as CASM^30^ and atg8ylation^20,79^. We have previously shown that, in *Drosophila*, obesogenic diets dysregulate phosphatidylethanolamine (PE) homeostasis^80^. Atg8/LC3 family proteins are conjugated to PE-lipids^18^, which makes the Atg8/LC3 proteins highly tuned to respond to membrane stress ^20^. Atg8 family members are recruited to lipid droplets to promote lipolysis on high-fat regimes ^13,14^, and autophagic pathways promote preadipocyte differentiation^15^. Hence, high-fat diet-induced membrane changes will likely influence Atg8/LC3-mediated leptin secretion.

In conclusion, Atg8-LC3 regulates nutrient surplus signals Upd2/leptin and is pivotal to the starvation response. Hence, we propose that Atg8-LC3 acts as a ‘bidirectional’ switch protein. By coupling scarcity and surplus mechanisms, Atg8-LC3 proteins enable organisms to manage nutrient flux effectively. In the future, decoding other bidirectional molecular switches and their mechanisms of action will enable us to develop new strategies to treat disorders that arise from nutrient imbalance states, such as obesity and anorexia.

## Acknowledgments

We thank Helene Knævelsrud for sharing the FLP-out Gal4 lines generated by Tom Neufeld. We thank David Strutt for sharing sequences for designing the V_H_H tagged constructs. The University of Washington undergraduate researchers Sunidhi Ranganathan, Jordan Wang, Hannah Goldfarb, and Vienna Wang provided us with technical assistance. We thank Phil Gafken and Lisa Nader in the proteomics shared resource at the Fred Hutch for assistance with IP-MS and ChenWei Lin for proteomic data analysis. We thank Alyssa Dawson and Katie Kim in the genomics shared resources at Fred Hutch for assistance with RNAseq and Pritha Chanana at Bioinformatics Shared resource for transcriptomic data analysis.

## Author contributions

Conceptualization: A.R., AEB, AM, KPK, MAA; Investigation: AM, KPK, MEP, CES, AEB, MAA, AR; Formal analysis, Data Curation: MEP, CES, KPK, AM, AEB, AR, J.D.; Visualization: J.D., AM, MEP, AEB, CES, AM, KPK; Supervision: A.R.; Writing-original draft: A.R.; Writing-review and editing: AM, KPK, MAA; Funding acquisition: AR.

## Data and materials availability

All data is available in the main text or the supplementary materials. Materials will be freely shared on request.

## METHODS

### Experimental Animals: *Drosophila melanogaster*

Only males were used in experiments at 5-10 days post-eclosion. Flies were cultured in a humidified incubator at 25°C with a 12h light-12h dark cycle and were fed a standard lab diet containing per liter: 15 g yeast, 8.6 g soy flour, 63 g corn flour, 5g agar, 5g malt, 74 mL corn syrup. For acute starvation, adult male flies, seven days old, were subjected to 4-hour starvation on 0% sucrose in agar at 29°C. For survival curves, flies were subjected to 1% sucrose in an agar diet. For RNAi experiments, flies were raised at 18°C until seven days post-eclosion, then shifted to 29°C for seven days.

A. The following fly strains used in this study were from our previous work ^2,38^ and obtained from other previously published work and the Bloomington *Drosophila* stock center (BDSC): *Upd2Δ3-62* (*Upd2Δ*; ^10^), *Lpp-Gal4* ^81^, UAS-Dicer; *r4-mCherry-Atg8a, Act5C>CD2>GAL4, UAS-GFPnls/TM6B* (kind gift from Helene Knævelsrud, University of Oslo) [Arsham and Neufeld, 2009^82^]. The control UAS strain used for over-expression experiments is UAS-Luciferase ^2^. RNAi lines from the TRiP facility at Harvard Medical School (http://www.flyrnai.org/TRiP-HOME.html) include *Luciferase-RNAi* (JF01801), Atg8-RNAi #1 (JF02895, BDSC 28989), Atg8-RNAi #2 (HMS01328, BDSC 34340).
B. The following UAS lines were generated for this study: *UAS-Upd2::GFP, UAS-Upd2-AIM-::GFP, UAS-VHH-Myc::Atg8-WT, UAS-VHH-Myc*. All transgenes of the same gene were inserted at the same att site to control expression levels—attp40 or attP2. For nanobody-based interaction experiments in Fig. 2A, flies of genotype *Lpp-Gal4*; *UAS-VHH-Myc::Atg8-WT (*experiment*) and Lpp-Gal4*; *UAS-VHH-Myc (Control)* were crossed to *UAS-Upd2-WT::GFP* or *UAS-Upd2-AIM::GFP*, the crosses were set at 25°C and the adult flies were dissected and stained at Day 7. For the clonal analysis experiments in Fig. 2D and Extended Data Fig. 2, virgin females of *Upd2-WT::HA; +; UAS-Luciferase-RNAi* and were crossed to males of yw hsflp; UAS-Dicer*; r4-mCherry-Atg8a, Act5C>CD2>GAL4, UAS-GFPnls/TM6B* (kind gift from Helene Knævelsrud, University of Oslo^82^). Crosses were set at 25C for 24-48 hours, parents were transferred to fresh vials, and vials with embryos were incubated at 29°C till eclosion. F1 flies were aged to 5-7 days at 29°C. Leaky expression at 29°C of the heat-shock inducible FLP leads to stochastic "flip out" activation of GAL4 expression ^83^ in a minority of fat body cells, activating UAS-driven expression of GFP and the inserted RNAi, allowing assessment of these cells in otherwise normal animals. Fat bodies were fixed and imaged as noted. *UAS RNAi* experiments and temporal over-expression in Fig. 2 flies were generated with the following genotype: *Upd2-crGFP; Ppl-Gal4, TubGal80ts* was crossed *to UAS- Atg8-RNAi, UAS-Luciferase-RNAi* for 2E; after eclosion at 18°C, adult male flies were maintained at 29°C for 7-10 days to inactivate the Gal80 and allow for Gal4 expression. Then 10-12 adult male flies were dissected and stained per genotype.
C. The following genomic tag-knockin lines into the endogenous Upd2 locus were generated for this study: *Upd2-WT::GFP, Upd2-WT::HA* and *Upd2-AIM::HA*. GFP or HA-tagged Upd2 CRISPR lines were developed in collaboration with WellGenetics Inc. using modified methods of Kondo and Ueda ^84^. To generate the HA-tagged Upd2 genomic knock-in used in Fig. 2B, the following strategy was used. In brief, the following gRNA sequence was cloned into U6 promoter plasmids for HA lines: TCCAATGAGTCTTGAGCCCT[CGG]/GCCGAGGGCTCAAGACTCAT[TGG]. Cassette 3xHA RFP containing 3xHA and 3xP3 RFP and two homology arms were cloned into pUC57 Kan as donor template for repair. Upd2/CG5988 targeting gRNAs and hs-Cas9 were supplied in DNA plasmids, together with donor plasmid for microinjection into embryos of control strain w1118. F1 flies carrying selection markers of 3xP3 RFP were further validated by genomic PCR and sequencing. CRISPR generates a break in Upd2/CG5988 and is replaced by cassette 3xHA RFP. 3XP3 RFP, which facilitates genetic screening, was flipped out by Cre recombinase. 46 bp) remains after excision between Stop codon and 3’UTR. To generate the *Upd2-AIM* point mutations, as seen in Fig. 2C, CRISPR/Cas9 mediated genome editing by homology dependent repair (HDR) using 1 guide RNA and a dsDNA plasmid donor was used. We introduced point mutations in the *Upd2-WT::HA* flies. To generate the AIM mutations, to replace W310 to A, TGG to GCG, and L313 to A, CTG to GCT of Upd2-WT::HA, using a targeting design without deletion. The following gRNA sequence was clones into U6 promoter plasmid: TATTGCGTCGCGGTCGCCGC[AGG]. Genomic PCR and sequencing methods to verify CRISPR alleles of Upd2/CG5988 by testing if the knock in cassette PBacDsRed is correctly used as template for homology directed repair and has been incorporated into guide RNA cutting site. For GFP line knockin line of Upd2-WT::GFP used in Fig. 2E gRNA sequence TCCAATGAGTCTTGAGCCCT[CGG] was used.

### Cell lines

*Drosophila* S2R+ cells, HEK293T cells and Human subcutaneous preadipocytes (Lonza Catalog #: PT-5020) were used for all cell culture-related experiments. ***Drosophila* S2R+ cells** were maintained in Schneider’s medium (GIBCO), 10% heat-inactivated FBS (SIGMA), and 5% Pen-Strep (GIBCO) at 25°C. For starvation, cells were incubated in Schneider’s Insect Medium without Amino Acids (United States Biological, Cat# S0100-01.10) for the indicated amount of time. **HEK293T cells** were cultured in DMEM with 10% FBS and 1% penicillin-streptavidin supplement. **Human Subcutaneous Preadipocytes** were utilized for human Adipocyte experiments. Preadipocyte cells were maintained in preadipocyte growth media-2 (Lonza Cat #PT-8002).

#### Cloning and Transgenic Flies

All cloning was done using the Gateway® Technology. For Atg8, entry cDNA was PCR amplified from Atg8 cDNA (clone LD05816 DGRC-Gold collection) and cloned into pENTR-D/TOPO using B.P. reaction (Gateway® BP Clonase II enzyme mix, Cat#11789-020, Invitrogen). For *Drosophila* Upd2 variants, pENTR-Upd2 ^38^ made for previous work in our lab was used. For site-directed mutagenesis of putative AIM sites, pENTR-Upd2 was mutagenized to convert tryptophan and leucine encoding codons to alanine. For human leptin ORF clone (Origene; pCMV-leptin-DDK-Myc; CAT#: RC209259) was used for mutagenizing the AIM sequences. Mutagenesis was done using the Q5® Site-Directed Mutagenesis Kit from NEB (Cat # E0554S). The sequence of oligonucleotides used for this mutagenesis reaction is available on request. The entry vectors were then moved using L.R. clonase reaction (Gateway® LR Clonase® II Enzyme mix, Cat#11791-020, Invitrogen) into destination vectors compatible with fly transformation, or cell culture with the appropriate C-terminal tags for Upd2 and N-terminal V_H_H tags for Atg8. Transgenic flies were generated by using the microinjection service provided by Bestgene Inc. The Myc tagged human leptin clones were subcloned to include a GFP tag for ELISA.

For generating the V_H_H tagged constructs, the sequence for V_H_H tag and appropriate linkers, utilized in a prior study by Strutt and colleagues ^45^, was appended to either Myc (control: V_H_H-Myc), Atg8-WT (V_H_H-Myc::Atg8-WT), or Atg8-PE- (V_H_H- Myc::Atg8-PE-). Double-stranded sequences for the three constructs (available on request) were synthesized using Codex DNA Inc. Synthetic DNA platform and cloned into PDONR221 using B.P. clonase (Gateway™ BP Clonase™ II Enzyme mix; Catalog number: 11789020, Invitrogen). Then, to generate expression clones, the PDONR vectors were cloned into expression competent S2 cell culture vectors from the Murphy Collection available from DGRC using L.R. clonase or into Gateway Vectors for fly transgenesis.

#### Triglyceride Measurements

TAG assays were carried out as previously described in (Rajan et al., 2017). In brief: Flies were homogenized in PBST (PBS + 0.1% Triton X-100) using 1mm zirconium beads (Cat#ZROB10, Next Advance) in a Bullet Blender® Tissue homogenizer (Model BBX24, Next Advance). Samples were heated to 70°C for 10 minutes, then centrifuged at 14,000 rpm (in a refrigerated tabletop centrifuge). 10.0 μl of the supernatant was applied to determine the level of TAG in the sample, using the following reagents obtained from Sigma: Free glycerol (cat # F6428-40ML), Triglyceride reagent (cat# T2449-10ML), and Glycerol standard (cat# G7793-5ML). Three adult males were employed per biological replicate. Note: For adult TAG assays, the most consistent results, with the lowest standard deviations, were obtained with 10-day old males. TAG readings from whole fly lysate (n=4 replicates of 3 flies each) were normalized to the number of flies per experiment. The normalized ratio from the control served as a baseline, and data are represented as fold change of experimental genotypes with respect to the control.

#### Feeding Assays

For hunger driven feeding analysis, age-matched adult male *Upd2-WT::HA* and *Upd2-AIM::HA* flies were given a normal diet for 7 days. 16 hours prior to feeding behavior assessment, half of the flies from each treatment were moved to starvation media (0% sucrose, agar). During the 3-hr assessment window of feeding behavior, individual flies were placed in a single well of FLIC (Fly Liquid-Food Interaction Counter) and supplied with a 5% sucrose liquid diet for all FLIC experiments. Detailed methods for how FLIC operate can be found in Ro et al., 2014^49^. All FLICs were performed at 10 am local time. For each experiment, half of the wells (n = 6/FLIC) contained the fed group, and the other half contained the starved group of flies for direct comparison. A total of 32–59 flies were measured for analysis of feeding from 4 independent experiments. Any signal above 40 (a.u.) was considered a feeding event. FLIC data and feeding events were processed using R. Graphing and statistical analysis (Normality test, Kruskal-Wallis Test, and Dunn’s multiple comparison test) were performed using GraphPad Prism 9.3.1.

#### Survival Assays

Survival curves were done using flies harvested in a 24-hour time frame and aged for 7 days. Ten males per vial were flipped onto 1% sucrose agar starvation food. The number of dead flies was recorded each day until every fly had died. Flies were kept in a 25°C incubator with a 12-hour light-dark cycle for the entirety of the experiment. Survival analysis was performed using the Survival Curve module of GraphPad Prism. A Mantel-cox test was the test used to determine statistical significance. 100-150 flies were used per genotype per curve. Data consolidated from 3 independent experiments is shown.

#### Drosophila adult adipose tissue preparation for fixed immunostaining

For fat body preparation, incisions using dissection scissors (Cat# 500086, World Precision Instruments Inc) were made to release the ventral abdomen from the rest of the fly body. Flies used for dissection were adult males, 7-10 days old. Dissections were done in Ringer’s medium (1.8 mM CaCl2, 2mM Kcl, 128mM NaCl, 4mM MgCl2. 35mM sucrose, 5mM HEPES). The tissue was fixed in 4% formaldehyde for 20 minutes and rinsed with PBS. For immunohistochemistry, the fixed fat tissue was permeabilized in PBS+ 1.0% Triton-X-100 for 3X washes 5 minutes, subsequently washed with PBS+0.3% Triton-X-100 (Fat wash). Blocked for 30 minutes at room temperature (R.T.) with gentle agitation in Fat wash+ 5% Normal donkey serum (Block). Primary antibodies, diluted in block, included Rabbit-anti-RFP (1:500, Rockland, #600-401-379); Chicken-anti-GFP (1:2000, Abcam, #ab13970); rat-anti-HA (1:500; Chromotek, #7C9); mouse-anti-Lamin (1:100; ADL67.10 DSHB) and incubated overnight at 4C. Washed 3X-5X for 15 mins each the following day in the fat wash at (R.T.) incubated with appropriate secondary antibody (from Jackson Immunoresearch) at a concentration of 1:500 in the block for 2-4 hours at RT. Washed 3X-5X for 5-15 minutes in the fat wash, mounted in SlowFade Gold antifade reagent with DAPI (Cat# S36938, Invitrogen). Images were captured with Zeiss LSM 800 confocal system and analyzed with Zeiss ZenLite or ImageJ.

***Drosophila* S2R+ cells** were passaged to 60-80% confluency. Cells were cultured in 96 well plates for transfections related to ELISA experiments and coimmunoprecipitations in 6-well plates. For ELISAs, cells were transfected with 20ng/well of pAc-Upd2::GFP 3 or indicated Upd2 variants, 10ng/well pACRenilla::Luciferase, and 150ng of dsRNA/well for knockdown experiments. For coimmunoprecipitations, cells were transfected with 200ng/well of the indicated plasmid.

For imaging experiments, cells were seeded on poly-D-lysine coated 8-well chambered culture slides for fixed imaging (MatTek CCS-8). The T7 flasks, which were 100% confluent, were used for seeing cells at 1:10 dilution at 400μl per well of the 8-well chamber dish. Transfections were done with plasmids indicated. For 8-chamber slides, 100ng/well of each plasmid was transfected. Transfections were done using the Effectene kit (Cat# 301427, QIAGEN) per the manufacturer’s instructions.

#### Drosophila S2R+ dsRNA production and cell treatments

Amplicons for dsRNAs were designed using the SnapDragon dsRNA design tool (https://www.flyrnai.org/snapdragon) and in vitro transcribed (IVT) using MEGAscript® T7 Transcription Kit (Cat# AMB1334-5, ThermoFisher). IVT reactions were carried out as per the protocol provided by the DRSC (available at: http://www.flyrnai.org/DRSC-PRS.html). The sequence of amplicons used in this study can be found in the table below. LacZ dsRNA was used as controls. All dsRNA knockdown experiments were carried out using two independent dsRNAs per gene. We found that this produced a knockdown efficiency of >85% (based on qPCR analysis) in S2R+ cells. S2R+ transfected with dsRNAs were incubated for four days to allow for gene knockdown. On the 4th day, the media was changed, and the ELISA assay was carried out on the 5th day. Note that the data is represented as percent change in Upd2/Leptin secretion normalized to transfection efficiency with 0% change indicating baseline secretion level.

See the ELISA assay procedure below.

#### Primer sequences used for dsRNA production

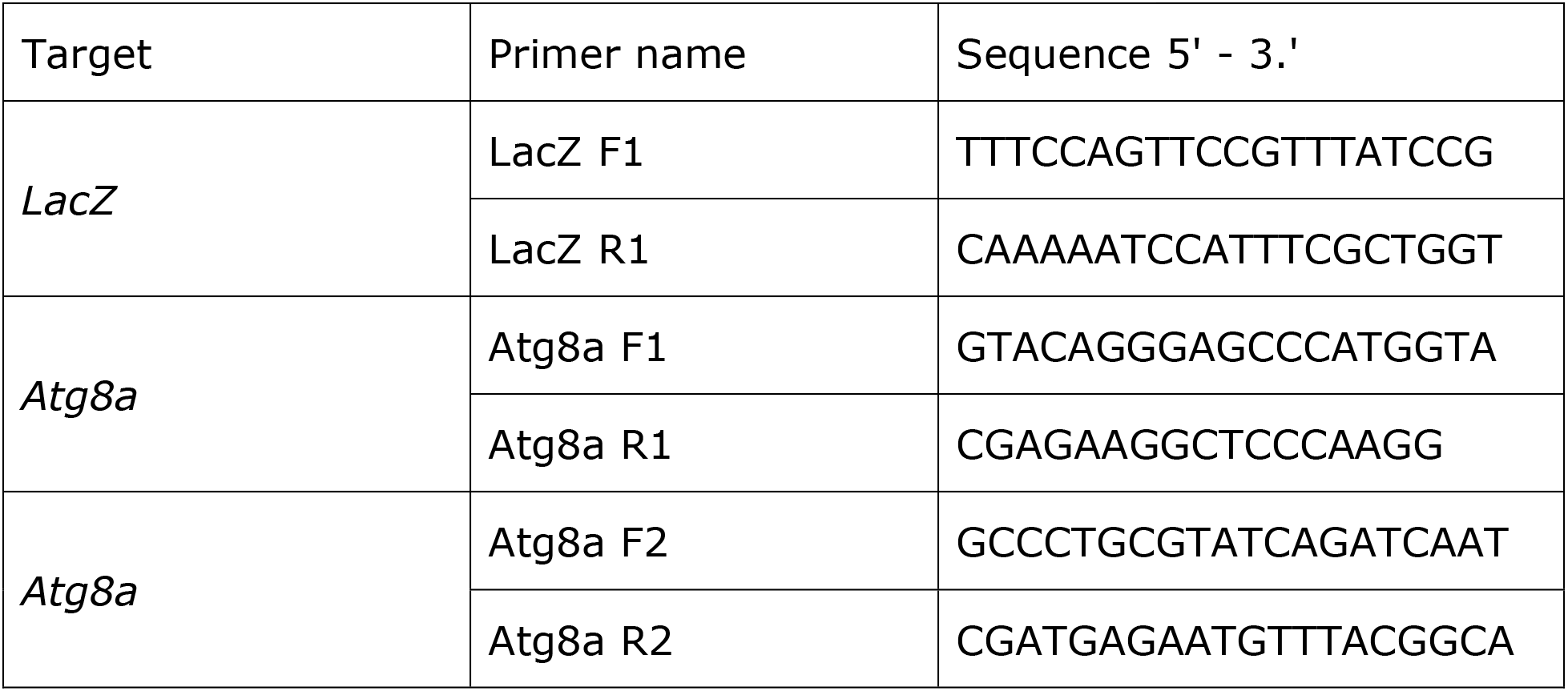

#### Treatment of cells with drugs

For drug treatment experiments, the media was replaced with media containing the drug on day 3 after transfection with Upd2::GFP. 4 hours later; the conditioned media was used for ELISA or imaging. Drugs used in this study include Torin1 (Cat# ab218606, Abcam). Stocks solutions of the drugs were made in DMSO as per the manufacturer’s instructions and used at a working concentration indicated in the Fig. 1C legends. DMSO treated replicates were used as controls. Note that for ELISAs, the data is represented as percent change in Upd2 secretion normalized to transfection efficiency, with 0% change indicating baseline secretion level.

**Human HEK293T cells** were cultured in DMEM with 10%FBS and 1% penicillin-streptavidin supplement. At the start of experiment, cells were trypsinized and seeded onto a 24 well plate or 8 well MatTek tissue culture slides for imaging (MatTek CCS-8) and grown to confluency and transfected with human Leptin constructs using Effectene kit (Cat# 301427, QIAGEN).

#### GFP sandwich ELISA assays

GFP sandwich ELISA assay was used for detecting Upd2::GFP and human Leptin::GFP. On day 1, 96 well medium bind polystyrene plates (Cat#CLS3368-100EA, Sigma) were incubated overnight at 4C with coating antibody (Cat# ACT-CM-GFPTRAP, Allele Biotechnology) diluted in 0.01M pH8.0 bicarbonate buffer at a concentration of 1μg/ml. On day 2, plates were washed briefly with PBS, blocked for 30 minutes with 1% BSA block in PBS, coated with conditioned media, and incubated overnight at 4C. Recombinant GFP protein (Cat# MB-0752, Vector labs), diluted in S2R+ cell growth media (64ng/μl to 4 ng/μl), was used in every ELISA plate as a positive control to ensure linearity of GFP readings. On day 3, the plates were washed with PBS+0.05% Tween-20 (PBS-T) blocked with 1% BSA in PBS for 30 minutes at RT. GFP detection antibody (Cat# 600-401-215, Rockland) was added to the diluted 1:1000 in 0.1% BSA in PBS-T. Plates were washed with PBS-T and incubated with secondary HRP conjugated anti Rabbit secondary antibodies (Cat# ab136636, Abcam) diluted at 1:5000 in 1% BSA block. Plates were washed in PBS-T with a final wash in PBS. For detection, each well was incubated 100μl 1-step Ultra-TMB ELISA substrate (Cat# 34028, Pierce), which was previously equilibrated to R.T., for approximately 5-15 minutes until detectable blue colorimetric reaction occurred. The reaction was stopped with 2N sulphuric acid, and absorbance was measured at 450nm. The TMB readings were normalized to transfection efficiency as measured from Renilla Luciferase assays (see below).

##### Renilla Luciferase assay

On day 2 of the ELISA assay, after the conditioned medium was transferred for use in ELISA assays, cells were re-suspended in 50μl of PBS and incubated with 50μl/well Renilla-Glo® Luciferase reagent (Cat# E2710, Promega) for 10 minutes and read using a multiwell luminometer.

##### Quantification for ELISA assays

For ELISA assays, the ELISA signal readings are normalized to transfection efficiency; the data is represented as percent fold change from control used as the baseline. Specifically, the ratio of TMB readings to Renilla Luciferase readings is calculated. This ratio from the control is used as a baseline, and the data is represented as the percent fold change of experimental conditions with respect to the control. A 2-tailed t-test quantified statistical significance on 6 biological replicates per condition. Error bars represent % S.D. (Standard Deviation). For TAG analysis, statistical significance was quantified by a 2-tailed t-test on 3-6 biological replicates per condition. Error bars indicate % S.D. (Standard Deviation). For qPCR analysis, normalized gene expression levels were analyzed using one-way ANOVA. Error bars represent the standard error of the mean (SEM).

#### Differentiation of human primary adipocytes

Human Subcutaneous Preadipocytes (Lonza Catalog #: PT-5020) were utilized for human Adipocyte experiments. Preadipocyte cells were maintained in preadipocyte growth media-2 (Lonza Cat #PT-8002). For experiments, 15,000 preadipocyte cells per well were seeded into a 24 well plate and grown to 90% confluency. Once confluent, cells were then switched to differentiation media (Lonza Cat #PT-8002) for seven days to allow for differentiation into mature primary adipocytes and development of lipid droplets. All cells were housed in a 37°C Incubator at 5% CO2 and 90% humidity.

#### siRNA knockdown in human primary adipocytes

Cell culture media of mature adipocytes was replaced 5 days prior to ELISA. For siRNA knockdown experiments, media was replaced with 500uL of Accell siRNA delivery media (Horizon Discovery, Dharmacon Accell siRNA delivery media Cat B-005000-500) with 0.5uM of each siRNA following manufacturer instructions. siRNAs for this study included: control (Accell non-targeting pool, 5nmol, Cat # D-001910-10-05), and Human Accell SMARTpools for: positive control Leptin siRNA (LEP (3952), Cat # E-011074-00-0005), LC3 (MAP1LC3A(84457) [Cat # E-013579-00-0005], MAP1LC3B(81631) [Cat# E-012846-00-0005], and MAP1LC3C(440738) [Cat # E-032399-00-0005], and GABARAP ((11337) [Cat# E-012368-00-0005], GABARAPL1(23710) [Cat# E-014715-00-0005], and GABARAPL2 (11345) [Cat# E-014715-00-0005]).

#### Endogenous Leptin ELISA

Supernatant was then collected and used to measure leptin concentration after 5 days of treatment with siRNA. ELISA’s for leptin were performed using the bio-techne Human Leptin Quantikine ELISA kit (DLP00). On the same day as the supernatant was collected, an adipored assay was performed with the remaining cells (Lonza Adipored Assay Reagent #PT-7009) for each sample used in the ELISA to confirm similar levels of adipocyte development and intracellular triglyceride was present amongst samples being compared. Adipocyte differentiation was quantified via intracellular triglyceride levels determined through the use of Lonza AdipoRed Assay Reagent. In brief, after supernatant for Leptin ELISA assay was collected, the remaining cells were then washed in PBS, followed by adding 1mL PBS mixed with 30µL AdipoRed. Cells were incubated for 15 minutes, followed by immediate fluorescent measurement using the SpectraMax i3 plate reader with an excitation of 485nm and emission of 572nm. For each ELISA assay, samples had two technical replicates at three different dilutions to confirm accurate concentration based on a standard curve of known Leptin concentrations.

#### Cell immunofluorescent staining

##### Drosophila S2R+ cells

For immunohistochemistry, cells were fixed for 20 minutes in 4% formaldehyde, washed in PBS for 5 quick changes, permeabilized in PBS+ 0.1% Triton-X-100 for 3X washes 5 minutes, subsequently washed with PBS+0.1% Triton-X-100 (Cell wash). Blocked for 30 minutes at room temperature (R.T.) with gentle agitation in Cell wash+ 5% Normal donkey serum (Block). Primary antibodies – Rabbit-anti-RFP (1:500, Rockland, #600-401-379); Chicken-anti-GFP (1:2000, Abcam, #ab13970); mouse-anti-Lamin (1:100; ADL67.10 DSHB) – were diluted in block incubated overnight at 4oC. Washed 3X-5X for 15 mins each was the following day in the fat wash at (R.T.) incubated with appropriate secondary antibody at a concentration of 1:500 (from Jackson Immunoresearch) in the block for 2-4 hours at RT. Washed 3X-5X for 5-15 minutes in cell wash, mounted with SlowFade Gold antifade reagent with DAPI (Cat# S36938, Invitrogen).

Images were captured with Zeiss LSM 800 or Zeiss Elyra 7 with Lattice SIM confocal systems and analyzed with Zeiss ZenLite or ImageJ 14.

##### Human HEK293T cells

Hek293T cells were trypsinized and seed onto an 8-well MatTek tissue culture slides for imaging (MatTek CCS-8) with rat tail collagen I (ThermoFisher Cat# A1048301). Rat collagen was used to allow the cells to adhere and spread on the glass slides. 1:80 dilution collagen:PBS. 200uL per MatTek well and incubated at 37C for at least two hours. MatTek was then either placed in the fridge for future use or liquid was removed and MatTek used immediately, after a quick wash and seedes with HEK293T cells. Upon reaching confluency, cells were transfected with either a Myc-tagged wild-type leptin or a Myc-tagged leptin with the AIM regions point mutated. Transfection was performed using the Effectene transfection reagent (Qiagen). Cells were incubated in the transfection media for 24 hours, then replaced with standard supplemented DMEM media for another 24 hours before the tissue was fixed. The tissue was fixed in 4% formaldehyde for 30 minutes and rinsed with PBS. HEK293T cells were then washed with PBS+0.3% Triton-X-100 (Cell wash) three times for 20 minutes each wash. Cells were blocked for 1 hour in Cell wash + 5% NDS (Block). Primary antibodies, diluted in block, included Rabbit-anti-GM130 (1:200, #ab52649); mouse-anti-myc (1:500, #ab32) and incubated overnight at 4C. Cells were then washed five times for 10 minutes each the following day in Cell wash at room temperature. Cells were again subject to block for 30 minutes. The appropriate secondary antibodies (Jackson Immunoresearch) and DAPI were diluted in block at a concentration of 1:500 for 2 hours. Afterward, cells were washed five times for 5-15 minutes in the Cell wash, mounted with Prolong Diamond Antifade (ThermoFischer #P36961). The HEK293T cell imaging was performed with Leica Stellaris 5 with deconvolution Lightning software.

#### Image acquisition, Morphometrics and image analysis

All cell biological experiments adult *Drosophila* adipose tissue, HEK293Tand S2R+ cells, were perform 3 independent times at least. For fat imaging 6-8 fat explants per experimental condition was used. For cells, at least 10 Z-stacks, with 5-10 cells per field of view was captured per experimental condition. The confocal settings for laser power and gain were maintained consistent per experiment across different genotypes or dietary conditions. To prevent bias, the person performing the starved versus fed imaging was blinded to the sample’s status. Excel or GraphPad Prism 7 software was used for data quantification and generation of graphs. Image analysis pipelines were built in MATLAB R2020b, and the associated scripts are available on request to the lead author. The following four types of analysis were performed in 3D to quantify:

1. *The nucleus-to-whole cell intensity ratio of Upd2 in S2R+ cells:* Since the S2R+ cells were transiently transfected we used a ratiometric quantification (Nuclear vs whole cell) to assess Upd2 intensity. To generate the 3D nuclear masks, Lamin or DAPI stacks were first maximally projected along the z-axis. After global thresholding, basic morphological operations, and watershed transform, the locations of the nuclear centroids in the x-y plane were used to scan the z-stacks and reconstruct the nuclear volume. The whole-cell volume was approximated from the nuclear volume by dilation.
2. *Drosophila adipocytes Upd2 levels:* After global thresholding, basic morphological operations, and watershed transform, the locations of the nuclear centroids in the x-y plane were used to scan the z-stacks and reconstruct each nuclear volume. The whole-cell volume was approximated from the nuclear volume by dilation. Before measuring the whole cell mean intensity of Upd2, intensities were corrected for signal attenuation along the depth of the tissue. In brief, the mean lamin pixel intensity was recorded at each z plane (when present), and the intensity decay was fitted with a single exponential after intensity normalization. The Upd2 intensity was then adjusted at each z-planes before calculating its mean intensity.
3. *Drosophila adipocytes clonal analysis:* The clonal volume was defined after thresholding and applying morphological filters to the GFP signal. Upd2 signal was adjusted with respect to tissue depth, and the mean Upd2 pixel intensity was then measured inside and outside of the clone volume.
4. *Leptin analysis in HEK293T cells for mean intensity, polarization and co-localization*: Nuclei were segmented in 3D from the DAPI stack channel using our own trained Stardist3D model, a neural network-based approach to detect star convex polyhedral shaped objects, such as nuclei (https://doi.org/10.48550/arXiv.1806.03535). To quantify the polarization of the leptin signal around the nucleus, its distribution in the equatorial region of the cell was considered. The equatorial region of each cell was defined by selecting the z-plane where the nucleus surface is the largest. The cell area was extrapolated from the nucleus area by tessellation, and the resulting cell was divided into 9 sectors with equal central angles. The integral intensity of the leptin signal was then computed for each sector. To compare the leptin distribution between cells, a circular shift was applied, such that the sector with the

Once these cellular compartments had been defined, they were used as masks to extract intensity values of the signal of interest and to compute either the mean pixel intensity in the nucleus, the ratio of the integral intensities between the nucleus and the whole cell, or between the combined vesicles and the whole cell.

Two-sided Wilcoxon rank-sum tests were used to assess the statistical significance of pairwise comparisons between experimental conditions.

We determined that our data-points were normally distributed, based on two measures: i) A GraphPad outlier test did not identify any outliers in our data; and ii) the majority of our data points for a particular condition were relatively similar to one other, with only a small standard error of mean or standard deviation.

#### Immunoprecipitation, Mass Spectrometry and Proteomic analysis

For Immunoprecipitation (I.P.) of human leptin from HEK293T cells, cells were transfected with GFP tagged Leptin-WT and Leptin-Myc. Protein for each condition was prepared by lysing 1 well of a 6-well dish, 4 days after transfection. 2mg/ml was used per I.P. experiments performed with camelid antibodies GFP-Trap Magnetic Agarose (Cat# gtma-20, Chromotek); as per the manufacturer’s protocol. Immunoprecipitates were electrophoresed approximately 1 to 2 centimeters into a SDS-PAGE gel.

Coomassie-stained gel bands were proteolytically digested with trypsin as described (in Cheung et al.2017; ^85^). De-salted peptides were analyzed by LC-MS/MS using an Orbitrap Eclipse with FAIMS (field asymmetric ion mobility spectrometry) mass spectrometer. Collected data were analyzed with Proteome Discoverer v2.5 to generate label-free quantification (LFQ) values and identified peptides were filtered to a 1% false discovery rate. LFQ data were normalized to the median value across all samples. Missing data were imputed with half of the global minimum value. P-values for pairwise comparisons were calculated by t-test. All plots were constructed by ggplot2 in R. Proteins that were predicted to interact with Leptin-WT exclusively as per the IP-MS experiment, were input into the STRING database (^58^. Kmeans clustering of the genes was conducted to categorize the genes into 3 clusters based on both known and predicted protein-protein interactions on the basis of numerous sources, including experimental repositories, computational prediction methods, and public text collections.

#### RNAseq

##### RNA prep

Total RNA prepared from 30 flies per genotype in triplicates, using the Direct-zol RNA miniprep kit (Zymo Research, cat#R2071). cDNA prepared with iScript cDNA Synthesis (Bio-Rad, cat#1708891), and 1mg RNA applied per reaction.

##### RNA quality control

Total RNA integrity was checked using an Agilent 4200 TapeStation (Agilent Technologies, Inc., Santa Clara, CA) and quantified using a Trinean DropSense96 spectrophotometer (Caliper Life Sciences, Hopkinton, MA).

##### RNA-seq Expression Analysis

RNA-seq libraries were prepared from total RNA using the TruSeq Stranded mRNA kit (Illumina, Inc., San Diego, CA, USA). Library size distribution was validated using an Agilent 4200 TapeStation (Agilent Technologies, Santa Clara, CA, USA). Additional library Q.C., blending of pooled indexed libraries, and cluster optimization were performed using Life Technologies’ Invitrogen Qubit® 2.0 Fluorometer (Life Technologies-Invitrogen, Carlsbad, CA, USA). RNA-seq libraries were pooled (35-plex) and clustered onto an S.P. flow cell. Sequencing was performed using an Illumina NovaSeq 6000 employing a paired-end, 50 base read length (PE50) sequencing strategy.

##### RNA-seq analysis

STAR v2.7.1^86^ with 2-pass mapping was used to align 50bp paired-end reads to *Drosophila* melanogaster genome build BDGP6.32 and Ensembl gene annotation 104 (http://uswest.ensembl.org/Drosophila_melanogaster/Info/Index). FastQC 0.11.9 (https://www.bioinformatics.babraham.ac.uk/projects/fastqc/) and RSeQC 4.0.0 ^87^ were used for Q.C., including insert fragment size, read quality, read duplication rates, gene body coverage, and read distribution in different genomic regions. FeatureCounts ^88^ in Subread 1.6.5 was used to quantify gene-level expression by strand-specific paired-end read counting. Bioconductor package edgeR 3.26.8 (https://academic.oup.com/bioinformatics/article/26/1/139/182458) was used to detect differential gene expression between conditions. Genes with low expression were excluded by requiring at least one count per million in at least N samples (N is equal to the number of samples in the smallest group). The filtered expression matrix was normalized by the TMM method ^89^ and subject to significance testing using a quasi-likelihood pipeline implemented in edgeR. Genes were deemed differentially expressed if absolute fold changes were above 2 and Benjamini-Hochberg adjusted p-values were less than 0.01.

---

**Figure S1.**
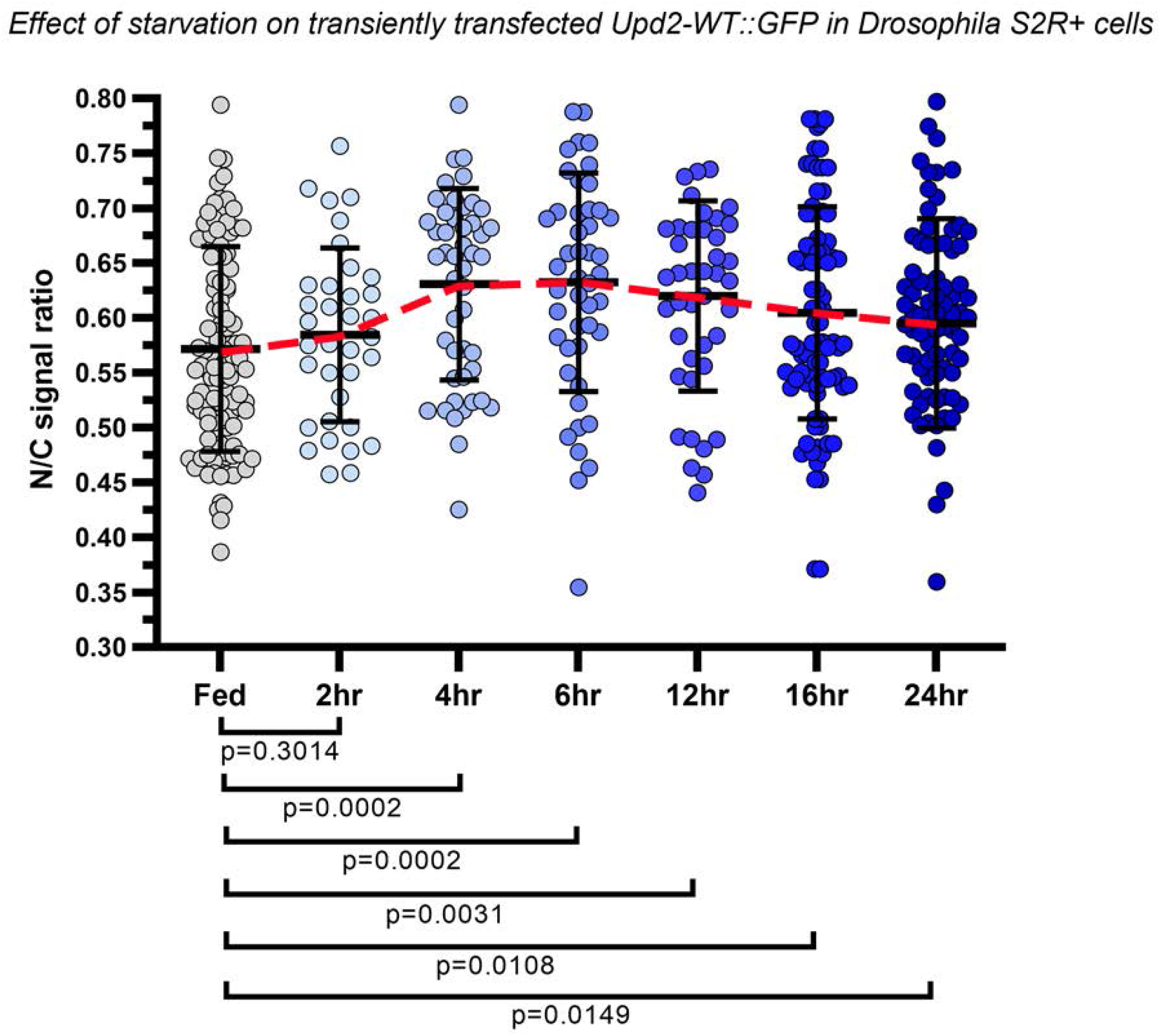
Companion to Figure 1. The ratio of Upd2::GFP nuclear/whole cell intensity is plotted over the time course (hours indicated on X-axis). Data is shown as mean ± SEM. Each dot represents a cell, 30-80 cells were counted per treatment condition. Statistical significance was measured by a non-parametric two-tailed Mann Whitney U test.

**Figure S2.**
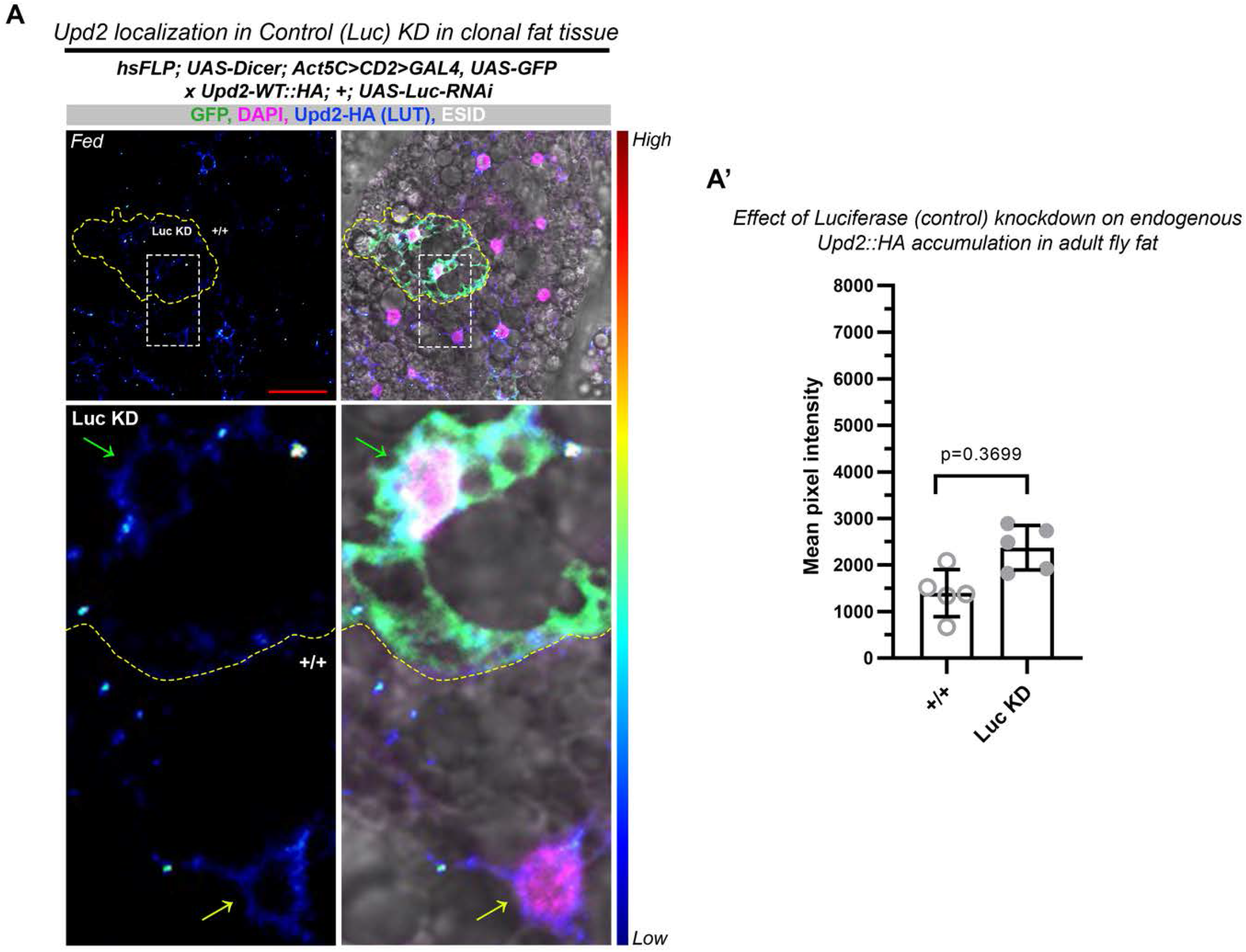
Companion to Figure 2. FLP-OUT technique was used to generate Act-GAL4-driven GFP-positive clones in adult fly fat in the background of the *Upd2-WT::HA* CRISPR knock-in. The effect of expressing control RNAi on HA-LUT signal in a positive clone (demarcated by a yellow dashed line), and bottom panel shows a higher magnification of the inset. The Upd2-HA-LUT signal within the positive knockdown of the clone relative to the surrounding wild-type tissue is quantified in A’. 5 clones were quantified to assess their significance.

**Figure S3.**
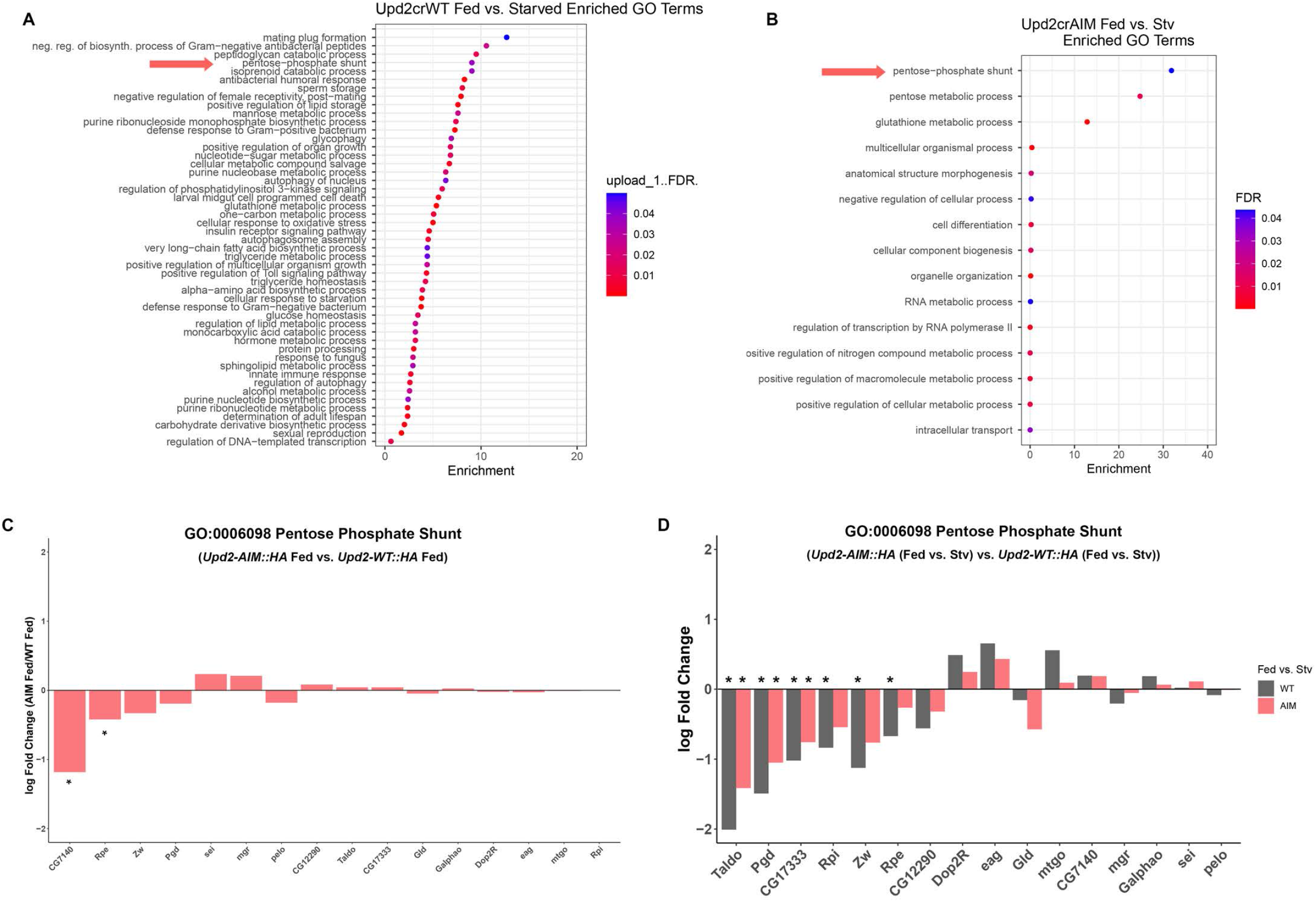
Companion to Figure 3. A, B. Gene Ontology (GO) plots are organized by highest enrichment, depicted on the X axis. False Discovery Rate (FDR) is indicated by the color of the dots (Blue to Red), with blue indicating an FDR=0.05 and red indicating an FDR<0.0001. C. Each bar represents the difference in log fold change between Upd2-WT and Upd2-AIM of the indicated gene in the x-axis. Differential gene expression was calculated using the edgeR Bioconducter package with significance calculated using Benjamini-Hochberg adjusted p-values (See Methods). Values of q<0.05 were considered significant. D. Each bar represents the difference in fold change between starved and fed state of the indicated genotype. Differential gene expression was calculated using the edgeR Bioconducter package with significance calculated using Benjamini-Hochberg adjusted p-values (See Methods). Values of q<0.05 were considered significant.

**Figure S4.**
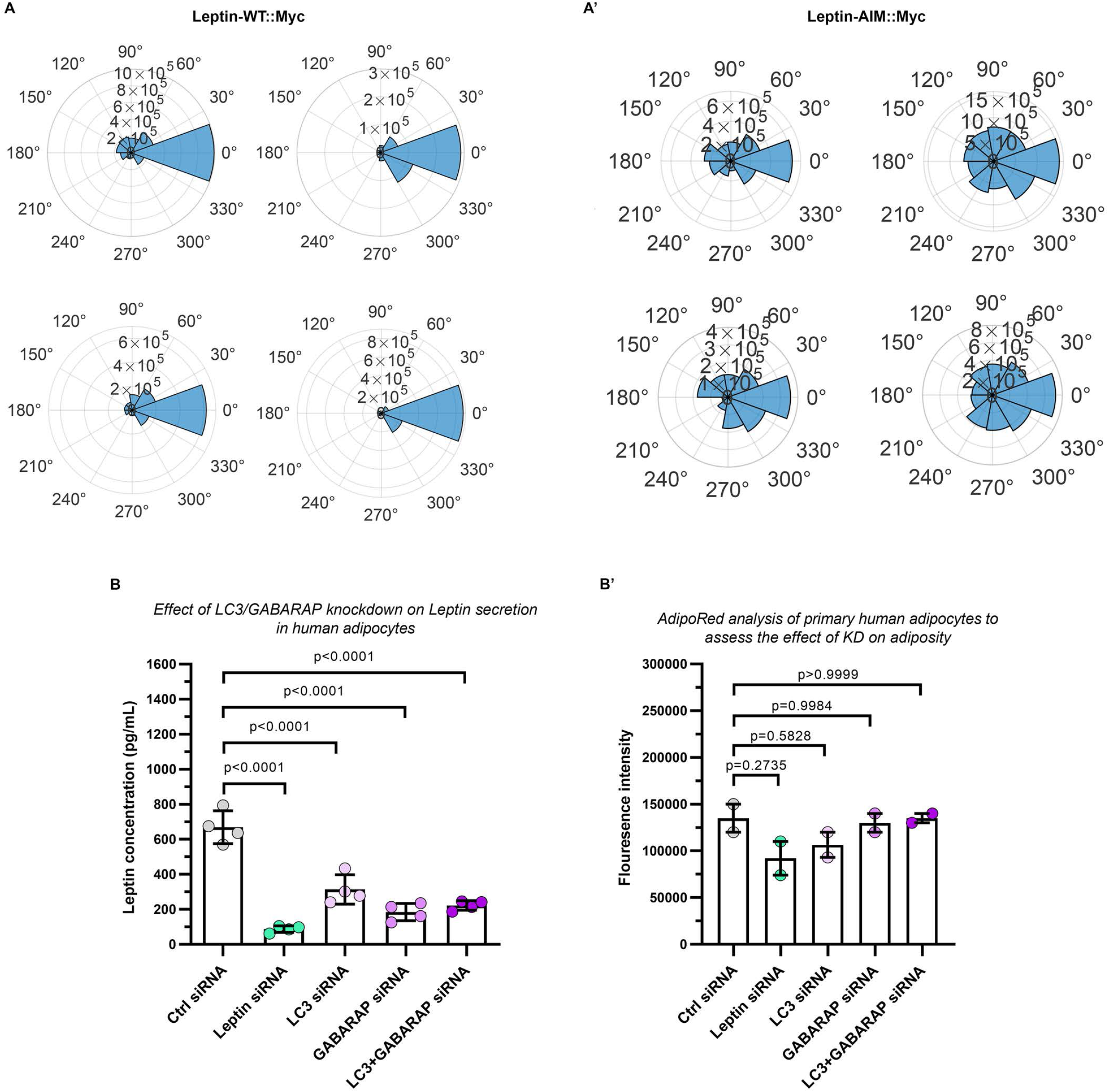
Companion to Figure 4. A, A’. Additional representative polarization plots for Leptin::Myc was calculated to describe and quantify its distribution. Leptin-WT::Myc (A’) shows a significantly higher polarization factor relative to Lepin-AIM::Myc. B. Endogenous Leptin ELISA from primary human adipocytes that were incubated with either non-targeted siRNA (positive control; grey), Leptin siRNA (negative control, green), LC3 siRNA (mauve), GABARAP siRNA (lilac), and LC3 + GABARAP siRNA (purple). Raw values of leptin are shown here, and see Figure 4E for % fold change. B’ AdipoRed staining of two independent siRNA treatments of adipocytes. Error bars represent % SD. Statistical significance was measured by a One-way ANOVA with Tukey corrections for multiple comparisons on 6 biological replicates per condition.

